# Ivermectin exposition during neurulation induces Neural tube defects and neuromuscular alterations in *Xenopus laevis* through purinergic P2X4-signaling

**DOI:** 10.64898/2026.06.19.733173

**Authors:** Claudio Catrupay-Valdebenito, Carlos F. Burgos, Bernardita Salgado-Martínez, Carla Vejar, Nicolás. A. Fuentes, Gonzalo E. Yévenes, Gustavo Moraga-Cid, Patricio A. Castro

## Abstract

**Background:** Neurulation is a fundamental process in the formation of the central nervous system (CNS). The process begins with the folding and fusion of the neural plate to form the neural tube which subsequently gives rise to the development of the brain and spinal cord. Environmental and genetic factors that disrupt neurulation can induce neural tube defects (NTDs) and consequently cause additional developmental complications, including motor impairments.

Purinergic signaling is a conserved form of extracellular communication (i.e. paracrine, synaptic signaling) that plays a role in early development. This signaling is mediated by purine nucleotides and nucleosides, which activate metabotropic P2Y and ionotropic P2X purinoceptors, respectively. Distinct patterns of intracellular calcium dynamics are observed throughout vertebrate development, from fertilization through organogenesis, including neurulation. Among P2X receptors, P2X4 is an ATP-modulated, Ca^2+^-permeable, ligand-gated ion channel characterized by having the highest Ca^2+^ permeability and is known to be modulated by ivermectin (IVM).

**Objective:** Our investigation focuses on assessing the effects of IVM treatment during neurulation and evaluating the impact of this drug on phenotype, motor behavior and neuromuscular junction (NMJ) structure at tadpole stage. These results were compared with those obtained following separate treatments with compounds that specifically block glycine, GABA(A) and nACh receptors, all which have been described as IVM targets.

**Results:** In this study we demonstrate the transcriptional expression for both P2X and P2Y purinergic receptors during neurulation, as well as the expression of P2X4. Following IVM neurula-treatments, we observed neural tube defects (NTDs), pigmentation changes, motor paralysis and alterations in neuromuscular junction (NMJ) structure, particularly affecting axonal branching. In contrast, treatment with the blockers strychnine, bicuculline and α-bungarotoxin, used to assess the involvement of GlyR, GABA(A)R and α7nAChR, respectively, failed to show similar outcomes.

**Conclusions:** In summary, our results highlight the critical role of purinergic signaling during early development, particularly P2X4 receptor mediated signaling during neurulation which may account for the pharmacological effects induced by the positive allosteric modulator ivermectin.

## Introduction

The formation of the central nervous system (CNS) is a crucial phase in chordates that begins with the folding of the epithelial neural plate to form the neural tube, process known as neurulation. Although this event exhibits some species-specific differences, it is highly conserved across vertebrates, including fish, amphibian and even mammalian species such as mice and humans (van der Spuy et al. 2023).

In the *Xenopus laevis* model, neurulation occurs between 14.5-21.5 hours post fertilization (hpf, stages 12-20; www.xenbase.org) whereas in humans it takes place between days 20-28 (Carnegie Stages, CS 9-13) (O’Rahilly and Müller 2007). Proper neural tube closure is regulated by spatiotemporal changes in cell shape and positioning, in which convergent extension, apical constriction and cell intercalation represent some of the most relevant cellular events involved in this process (van der Spuy et al. 2023). These cellular dynamics are guided by classical signaling pathways such as Wnts (Wingless-int), BMPs (Bone Morphogenetic proteins) and FGFs (Fibroblast Growth Factors) which collectively establish the initial framework of the nervous system that will give rise to the brain, brainstem, spinal cord and the diverse cellular populations of the nervous system (Steventon et al. 2009; Ten Donkelaar et al. 2006). Disruption of this process leads to the generation of neural tube defects (NTDs), the most common congenital malformation of the central nervous system worldwide, with an estimated prevalence of 1 per 1000 births in America (Ten Donkelaar et al. 2006; Zaganjor et al. 2016).

Besides the classical signaling pathways described above, neurotransmitters (NTs) participate in complementary and essential embryonic events, such as cell differentiation and migration that contribute to the proper formation of the CNS (Messenger et al. 1999; Moiseiwitsch 2000). In *Xenopus,* several classical neurotransmitter pathways have been implicated in normal embryonic development, including glutamatergic, adrenergic and dopaminergic signaling, which directly influence pigmentation and craniofacial structure development, among other structures (Sullivan and Levin 2016). Additionally, the involvement of NMDA receptors, the glutamate-synthesizing enzyme glutaminase (GLS) and the vesicular glutamate transporter (VGluT) during neurulation has been reported, supporting a role for glutamatergic signaling to an appropriate neural tube formation in mouse and frog (Sequerra et al. 2018; Tiboni et al. 2021; Goyal et al. 2026; Benavides-Rivas et al. 2020). However, the specific mechanisms underlying neurotransmitter participation during neurulation remain poorly understood.

Additionally, to NTs described previously, ATP, the principal purinergic signaling agonist, acts as a neuromodulator involved in several physiological processes including short-and long-term (LTP) synaptic communication as demonstrated in a P2X4 knockout mouse model. Hence, developmental alterations in *Xenopus*, such as NTDs and defects in neuromuscular junction (NMJ) have been associated to ATP signaling and P2 receptors (Fu 1995; Phillis 2024; Sim et al. 2006; Tovar et al. 2023).

Purinergic signaling comprises ionotropic P2X_1-7_ and metabotropic P2Y_1,2,4,6,11-14_ receptors along with ectonucleoside triphosphate disphophohydrolases enzymes (E-NTPDases). This signaling system is involved in neurulation and posterior neural development, where it regulates progenitor cell proliferation as well as neuronal and glial maturation and differentiation (Andrejew et al. 2023; Zimmermann 2011). Purinergic signaling is strongly involved and associated with calcium (Ca^2+^) dynamics via the activation of P2Y1-like Gα_q-_containing receptors (P2Y_1,2,4,6,11_), and is further complemented by Na^+^ and Ca^2+^ permeable P2X_1,2,3,4,5,7_ channel receptors, modulating processes ranging from neuronal induction to neural differentiation (Burnstock and Ulrich 2011; Von Kügelgen 2011; Webb and Miller 2003). Distinct patterns of intracellular Ca^2+^ dynamics, known as Ca^2+^i transients, are observed throughout vertebrate development, from fertilization to organogenesis (Sequerra et al. 2018; Slusarski and Pelegri 2007; Whitaker 2006; Goyal et al. 2026). Ca^2+^ fluxes play a crucial role in neurulation, regulating biomechanical processes such as apical constriction, cell polarization, cell intercalation, morphogenetic movement, and cell shape changes. Several studies have shown that dysregulation of Ca^2+^i signaling results in severe conditions like NTDs (Paudel et al. 2018; Sequerra et al. 2018; Suzuki et al. 2017; Goyal et al. 2026).

Cellular communication during early neural development, including neurulation, involves paracrine signaling in which the extracellular release of molecules such as ATP, glutamate, and ADP be mediated by connexins hemichannels (Cx46) (Tovar et al. 2023). Several examples of this type of signaling have been investigated, including the activation of P2X_7_R via autocrine and paracrine messengers that regulate early embryonic differentiation and cell fate through apoptosis of neural progenitor cells (Delarasse et al. 2009; Rodrigues et al. 2019; Suyama et al. 2012). These findings suggest an active role for intracellular Ca^2+^i mobilization mediated by paracrine purinergic signaling during development. However, the specific roles of individual purinergic receptors during early events of neural development, such as neurulation, remain incompletely understood.

P2X receptors are channels formed by either three homomeric P2X_1,2,3,4,5,7_ or heteromeric P2X_1/2-6, 2/3,5,6, 3/5, 4/5-7, 5/6_ assemblies of subunits. These channels are activated by ATP and permeable to Ca^2+^, Na^+^ and K^+^ ions, inducing rapid changes in membrane potential upon activation (Rodrigues et al. 2019; Khakh and North 2012; Saul et al. 2013). Among the P2X receptor family, P2X4 exhibits the highest Ca^2+^ permeability (Fractional Ca^2+^ current, Pf% of 16%), and is expressed in a wide variety of cell types, including central neurons and immune cells (Stokes et al. 2017; C. Shen et al. 2023; Samways et al. 2014).

Ivermectin (IVM) is a widely used antiparasitic drug considered safe due to its minimal side effects and selective action on nematode glutamate-gated chloride channels (GluCl). However, despite its safety profile, evidence of IVM toxicity in animals has been reported involving interactions on GABA_A_(GABA(A)R), glycine (GlyR) and alpha 7 nicotinic acetylcholine (α7nAChR) receptors, leading to hyperlocomotion and deleterious effects in several organs (Bates 2020; GabAllh et al. 2017; Krause et al. 1998; Krůšek and Zemková 1994; Löscher 2023; Parisi et al. 2019). Additionally, IVM is the strongest positive allosteric modulator (PAM) of P2X4 (exerting its effects at concentrations as low as 250 nM, while exhibiting minor or no effects toward other homomomeric P2X receptors) increasing Ca^+2^ conductance, potentiating the current and delaying the channel deactivation of P2X4R (Stokes et al. 2017; CRUMP and ŌMURA 2011; Khakh et al. 1999).

Furthermore, IVM has been proposed as an adjuvant therapy for refractory epilepsy, with some studies reporting a significant reduction in seizure frequency in affected patients. However, more recent studies have raised concerns regarding the safety of IVM for this therapeutic application (Diazgranados-Sanchez et al. 2017; Dusabimana et al. 2021; Mandro et al. 2020; Siewe Fodjo et al. 2019). Moreover, additional analyses have highlighted the lack of sufficient evidence to establish a conclusive safety profile for IVM, especially during pregnancy in women with epilepsy and throughout early development, where purinergic signaling is highly active (Löscher 2023; Nicolas et al. 2020; Tovar et al. 2023).

Based on these antecedents, we aimed to evaluate the role of P2X4-ATP signaling specifically during neurulation in *Xenopus* embryos using the antiepileptic and antiparasitic drug IVM. Using molecular approaches, we detected transcripts of P2X and P2Y subunits, and protein expression of P2X4R. Complementary bioinformatic analyses validated our *Xenopus laevis* model including the strong positive modulation of IVM on P2X4. We observed concentration-dependent effects on pigmentation in tadpoles and induction of NTDs which were associated with increased axonal branching and thus reduced synaptic contact, thereby explaining the impaired motor behavior and responses. Finally, no evident IVM-similar effects were observable blocking other possible targets such as GlyR, GABA(A)R and nAChR.

## Methods

### Ethical Statement and Experimental Animals

The care and handling of the *Xenopus* animals were approved by the ethics, bioethics and biosafety committee from Universidad of Concepción (CEBB 1459-2023, 30, May 2023). The animals were treated following the ethical protocols established by the United States National Institute of Health (NIH) and the amphibian euthanasia protocol was based on that described by (Close et al. 1997). Adult animals are euthanized by an overdose of tricaine methanesulfonate (MS222) anesthetic followed by cervical dislocation. A container is filled with 1 L of tap water and 50 mL of tricaine methanesulfonate stock solution (10 g/l MS222, buffered with sodium bicarbonate to pH 7.4) is added (final concentration: 0.5 g/L). The frog is placed in the container for at least ten minutes after frog appears lifeless. The euthanasia procedure is finalized by cervical dislocation. Male frogs’ belly is opened and testes removed. Testes are blotted on paper towel to eliminate excess of blood. Carcasses are kept in the freezer for later disposal. Testes are stored at 4°C in 1× MMR medium supplemented with 20% FBS and 1% gentamicin for up to 10 days. Tadpoles used for immunohistochemistry are euthanized on a Petri dish by MS222 overdose (final concentration: 0.5 g/L) and then fixed.

### Obtaining Xenopus laevis Embryos

*Xenopus laevis* oocytes were collected using the *in vitro* fertilization technique. Ovulation of the female frog was previously induced by subcutaneous injection of the Human Chorionic Gonadotropin hormone (hCG). 48 hours before fertilization (pre-prime), the female was injected with 50 units of hCG. 24 h later the female was injected again with 450–500 units of hCG (prime). Embryos were maintained in Marc’s modified Ringer’s solution (10% MMR, pH 7.4), containing 1 M NaCl, 20 mM KCl, 10 mM MgSO_4_, 20 mM CaCl_2_, and 50 mM HEPES. The male gonads were obtained through abdominal dissection. The testes were stored in 1× MMR medium supplemented with 20% FBS. 12 h after the second injection of hCG, the oocytes were collected in 1× MMR solution, later they were fertilized with *X. laevis* testis extract and incubated for 1 h at room temperature in 10% MMR solution. Afterward, the medium was replaced by a 2% cysteine solution in 10% MMR, pH 7.8–7.9 for 5 min, to dejelly embryos. Subsequently, the embryos were washed and incubated in 10% MMR at room temperature and finally harvested at different neurulation stages: 12.5 (early neurulation), 14 (middle neurulation), and stage 20 (late neurulation); and stage 40–45 (tadpole) (Nieuwkoop et al. 1994). We observed the expression of purinergic receptors in stages 12.5–20 and the analysis of the closure of the neural tube and morphological effects in stages 40–45. The adult brain as a positive control to validate purinergic receptors expression in molecular biology assays.

### Real-Time RT-qPCR

#### Total RNA Extraction

For the total RNA extraction of the embryonic stages of *X. laevis*: Stage 12.5, stage 14, stage 20, and adult brain tissue, the EZNA^®^ HP Total RNA kit was used, homogenizing the samples in 700 μL of the buffer lysis GTC. This volume was implemented according to the manufacturer’s instructions per amount of tissue (30 mg) (15 embryos). The samples were subjected to centrifugation and washed with 700 μL of 70% *v/v* ethanol according to the Kit instructions. Finally, the total RNA was resuspended in 30 μL of RNase-free water and quantified by measuring its absorbance at 260 nm and its purity according to the 260/280 ratio in the NanoQuant infinite 200 PRO equipment (Tecan, Zurich, Switzerland). RNA was stored at −80 °C for later use.

#### Reverse Transcription (RT) of Total RNA

DNA synthesis was performed using the reverse transcriptase enzyme M-MuLV Enzyme Mix (ProtoScript^®^ II First Strand cDNA Synthesis Kit, New England BioLabs, Ipswich, MA, USA), adding 1 μg of total RNA sample. For a final volume of 20 μL, the RNA was incubated with 0.5 μg of oligo-dT, denatured at 70 °C for 5 min, and placed on ice for 2 min. Subsequently, 400 U of the reverse transcriptase enzyme M-MuLV Enzyme Mix was added and incubated for 1 h at 42 °C and finally at 70 °C for 5 min.

#### Quantitative Real-Time RT-PCR (qRT-PCR)

The real-time RT-qPCR reaction was prepared with the Brilliant II SYBR Green qPCR Master Mix kit (Agilent Technologies, Santa Clara, CA, USA) in a final volume of 20 μL, containing 5 μL of diluted 1:10 cDNA and 500 nM of each splitter set (**Table 1**). Each reaction mixture was incubated at 95 °C for 10 min, followed by 40 cycles at 95 °C for 15 s, 60 °C for 15 s, 72 °C for 15 s in an AriaMx real time PCR system (Agilent Technologies, Santa Clara, CA, USA). The relative expression of purinergic was calculated by the ΔCT method using sub-1 gene as housekeeping. Subsequently, the ΔΔCt was determined by normalizing the conditions to stage 12.5. The negative controls for the amplification of the samples were treated with the same amplification protocol, but without adding cDNA to the mixture.

**Table 1.**
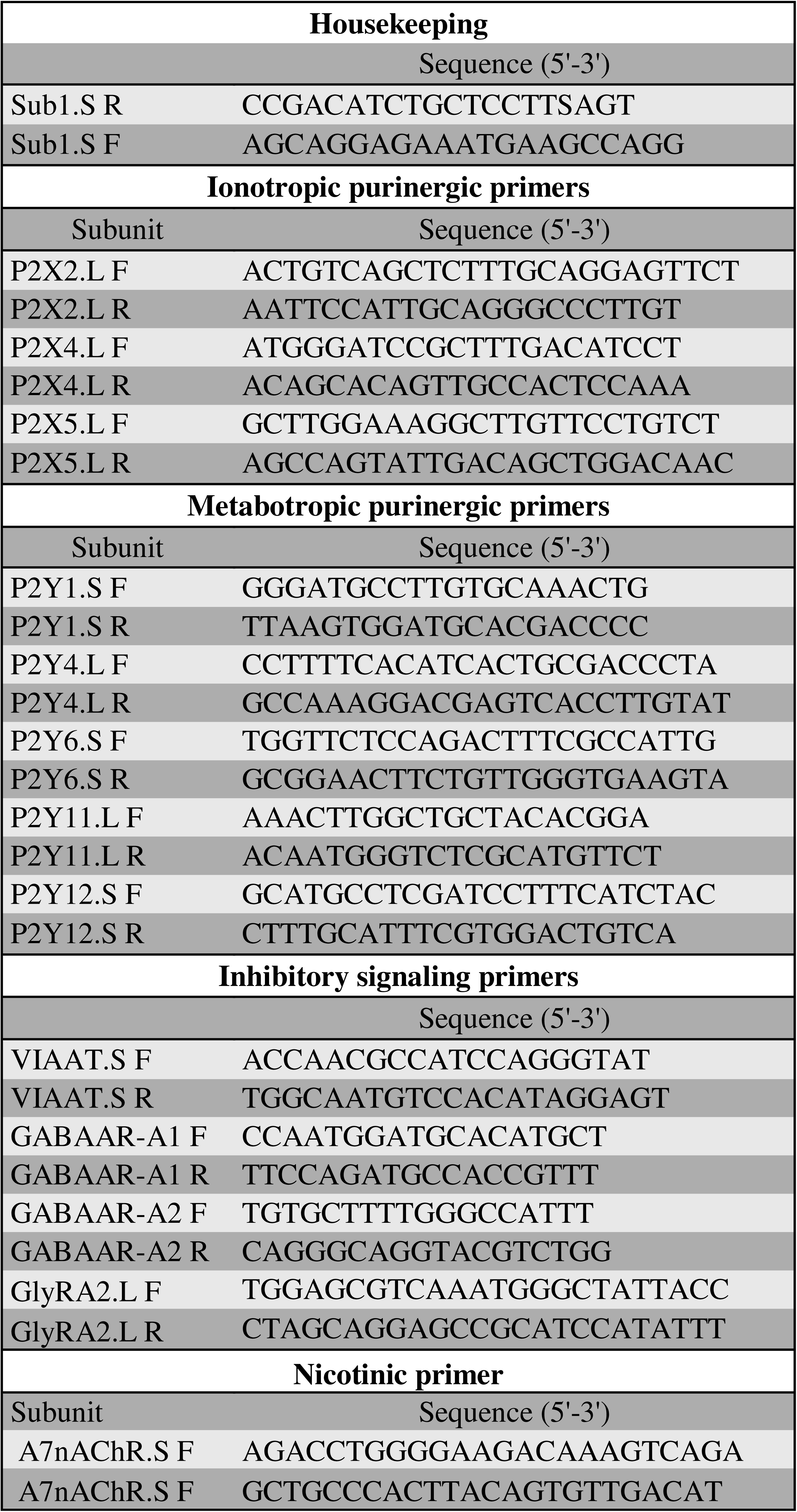
Primer sequences used in RT-qPCR and PCR neurula embryos. R=Reverse primer, F=Forward primer, S=Short, L=Long.

#### Pharmacological Modulation of the Neurulation Process

Stage 12.5 embryos were incubated with IVM (Cayman Chemical, USA), Stry, BIC (Tocris, UK) and α-BTX (Invitrogen) at 0,1; 0,5; 1, 3, 10, 30, 100 μM and MMR with DMSO 0,1; 0,3; 1% as a control. After 7 h of incubation, the embryos (stages 19–20) were washed three times with 10% MMR and kept in this solution until the tadpole stage (stage 40–45). Morphological analyses and image acquisition were performed using a SMZ25 stereo microscope, NIKON (Japan) coupled to a brightfield AmScope (MU300) camera, 1× objective (embryos stage 20) and 0.63× objective (tadpole embryos) and 2× objective (interocular tadpole region) in which the morphological alterations during the neurulation process were evaluated (Figure 1).

**Figure 1.**
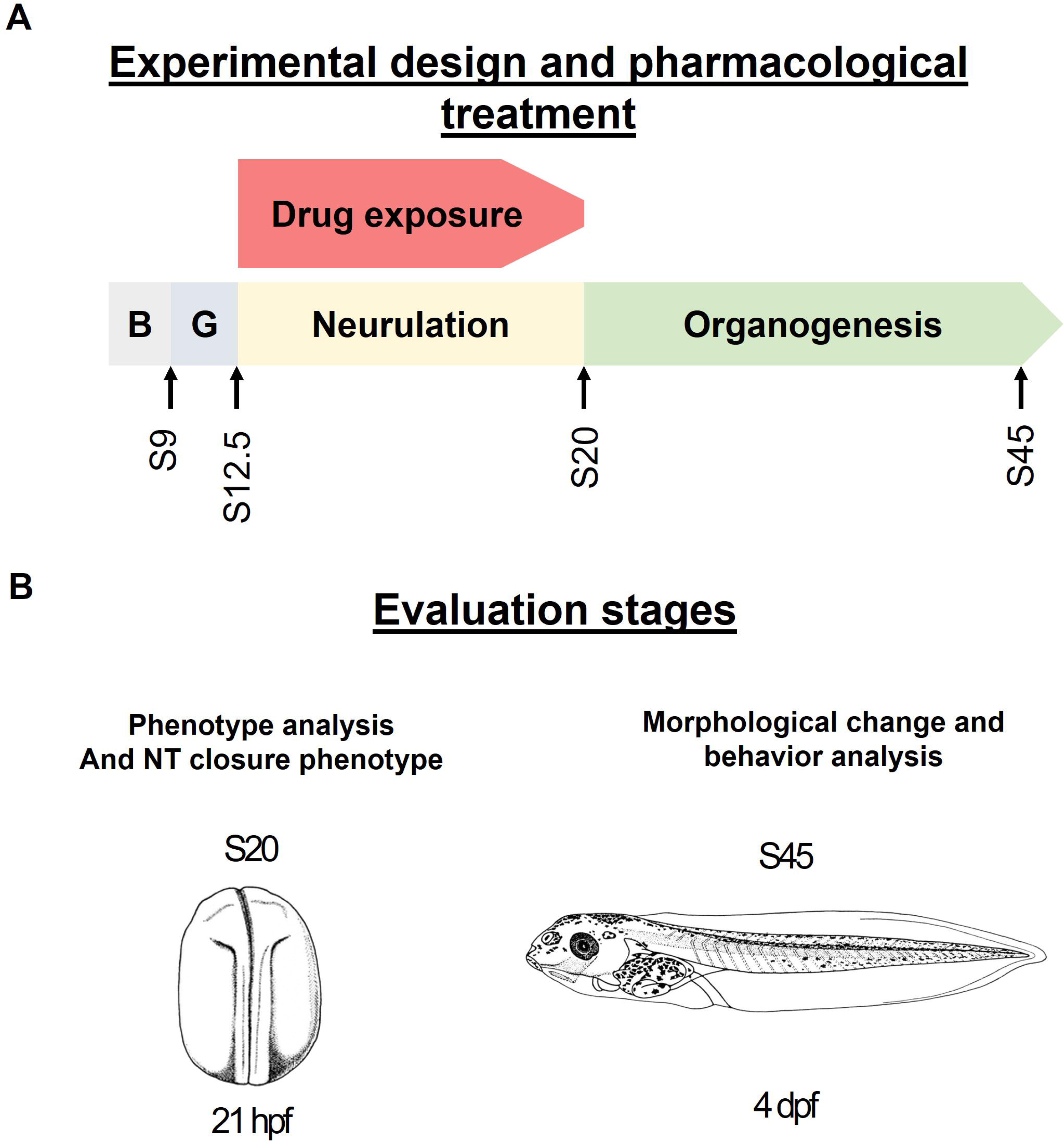
Diagram of pharmacological treatment and evaluation stages. **(A)** Scheme of pharmacological treatment during neurulation in *Xenopus laevis* development. **(B)** Evaluation stages at the end of pharmacological treatment in S20 and during organogenesis at S45. B=Blastula, G=Gastrula, S=Stage, NT= Neural tube. Hpf/dpf= Hours/days post fertilization.

#### Sensorimotor behavior

Tadpoles were positioned in the center of a well of a 12 well culture dish filled with 10% MMR. A hand-held metal tip (Fine science tools, USA) was used to lightly stroke the tadpole on the mid-tail of the resting tadpole, or until a swimming response was elicited. Responses were recorded on video at 30 Hz. Each tadpole was tested at least 3 times during thirty seconds. Stage 45 (S45) tadpoles were evaluated and categorized by swimming away (Normal), short swimming away (Reduced), or no response as described by Roberts and colleagues (Boothby and Roberts 1995; Roberts 2010).

### Western Blot Assay

#### Total Protein Extract

*Xenopus laevis* embryos in neurulation stages were used in this essay. Total protein extracts were manually homogenized (20 Gauge Needle/Syringe) using protease inhibitor cOmplete™ (ROCHE, Switzerland). The concentration of total lysed protein was determined using a Micro BSA kit (Thermofisher, Waltham, MA, USA) and NOVOstar equipment reading at 280 nm (BMG Labtech, Ortenberg, Germany).

#### Polyacrylamide Gel Electrophoresis and Electrotransfer

Proteins of interest were separated using a 12% denaturing acrylamide gel (SDS-PAGE). Samples were incubated at 95°C for 5 min in loading buffer (62.5 mM Tris-HCl pH 6.8, 2% SDS, 10% glycerol, 0.01% bromophenol blue) and 100 mM DTT. 240 µg of total proteins were loaded on the gel next to the protein standard (Spectra Multicolor, Broad Range Protein Ladder, Thermo Scientific, MA, USA) and ran at 100 V in run solution (25 mM Tris, 250 mM glycine, and 0.1% SDS). Proteins were transferred to Immobilon-P membrane (0.45 mm pore, Merck Millipore, Tullagreen, Carrigtwohill, Irland) with transfer solution (25 mM Tris, 192 mM glycine, 20% methanol) at 250 mA during 2 h. Power supplied by PowerPac™ Basic Power Supply and performed in a Mini Trans-Blot® Cell (2025 Bio-Rad Laboratories, Inc. United States)

#### Protein Immunodetection

Membranes containing the proteins were washed with TBS-Tween (150 mM NaCl, 10 mM Tris, 0.05% Tween20). Then blocked with 5% milk in TBS-Tween for 1 h. Overnight incubation at 4 °C using primary antibodies was carried out. The primary antibody was anti-P2X4, with an antigen sequence: KYKYVEDYEQGLSGEMNQ (1:250, Alome Labs, Jerusalem, Israel). The secondary antibody incubation was performed for 2 h using anti-rabbit peroxidase-conjugated antibodies (1:5000; ThermoFisher, Massachusetts, USA). The membrane was washed with TBS-Tween for 10 min five times. Finally, the membrane was revealed with a chemiluminescent solution (Western Lightening Plus-ECL, Perkin Elmer, Waltham, MA, USA) and the signal recorded at the chemiluminescent and fluorescent equipment (Odyssey FC, Li-COR, Lincoln, NE, USA).

Signal intensity measurements corresponded to the dynamic range over the background signal. Only a clear and distinguishable peaks relative to background signal were considered for analysis.

### Bioinformatic Analysis for P2X4-IVM Interaction

#### Modeling and Preparation of P2X4

We used the structure of P2X4 as template for comparative modeling to *Xenopus laevis* receptor from the Protein Data Bank PDB (4DW1), 2.8 Å resolution (Hattori and Gouaux 2012). The molecular compound structure of IVM (CID: 6321424) (MW: 875.1 g/mol) was extracted from the PubChem. Charges, protonation states, and rotatable bonds were assigned in the AutoDock Tools version 1.5.7 program (Trott and Olson 2010), before protein-ligand docking simulations.

The protein was modeled using the P2X4 amino acid sequence in *Xenopus laevis* (NP_001082067.1) obtained from National Center of Biotechnology Information (NCBI), and the P2X4 structure from zebrafish (PDB 4DW1) as template, due to its high sequence identity (59%) and hydrophobic amino acid conservation (77%), utilizing Modeller 10.4 (Eswar et al. 2006). Modeller generated 1000 initial models, the best of which were selected based on the ranking determined by an internal scoring function (molpdf) and DOPE (M. Shen and Sali 2006).

Lateral chains and missing hydrogens were added and disulfide bridges built at pH 7±0,2 through the Protein Preparation Wizard module from Maestro (Schrödinger, LLC, New York, NY, 2024). Energy minimization of the model was subsequently carried out using Macromodel (Schrödinger, LLC, New York, NY, 2024).

#### Protein-Ligand Docking and analysis

The XlP2X4-IVM interaction analysis was performed in the AutoDock Vina program (Eberhardt et al. 2021). The interaction parameters were made to keep the protein rigid and the ligands as flexible molecules. In addition to this, a Grid Box: (Coordinates X, Y, Z:-8.16, 8.275, 102.953) of size (X, Y, Z in Å: 48, 49, 36) on the transmembrane region (TM) was generated using the AutoDock tools program. The generated conformations were classified according to the affinity constants predicted in kcal/mol.

With molecular docking results we identify two pockets. This means that the ligand adopts different positions on the protein, which gives an approximation in terms of spatial orientation and pattern of protein interaction. Depending on production of affinity (A lower negative value) between IVM and P2X4, there exists a higher probability that produces a favorable and stable interaction (Nguyen et al. 2020).

We selected the conformation with highest affinity of two pocket were analyzed with LigPlot (Wallace et al. 1995) and interactive atoms involved in Discovery Studio software v.25.1.0.24284 (Dassault Systèmes 2025). The characterization of the protein-ligand interaction shows the different interactions detected in the protein-ligand conformation obtained.

#### Molecular Dynamics Simulations

Molecular dynamics (MD) simulations were performed using the Desmond (Schrödinger, New York, NY, 2025). The protein–ligand complexes for each binding site, previously identified via molecular docking, were selected based on their binding affinities. These complexes were embedded into a hydrated lipid bilayer model and simulated under physiologically relevant conditions. Specifically, the systems were integrated into a 1-palmitoyl-2-oleoyl-sn-glycero-3-phosphocholine (POPC) membrane and solvated using the SPC water model. To ensure electrostatic neutrality, a 0.15 M NaCl concentration was maintained. The system underwent a relaxation protocol comprising a 20 ns restrained MD phase, wherein harmonic positional constraints of 10 kcal/mol·Å² were applied to the protein backbone and ligand heavy atoms. Subsequently, all restraints were removed, and 100 ns production simulations were conducted under semi-isotropic NPT ensemble conditions. The temperature was maintained at 300 K using the Nosé–Hoover chain thermostat, and the pressure was regulated at 1.01325 bar through the Martyna–Tobias–Klein barostat. The OPLS4 force field was employed throughout the simulation (Millar-Obreque et al. 2025). Trajectory data were collected over the entire simulation period, and quantitative analyses, including Root Mean Square Deviation (RMSD) and residue contact profiles, were performed.

#### Whole mount Immunofluorescence staining

To analyze morphological changes in the NMJ, we labeled the cellular components of the NMJ: Axon motoneurons and acetylcholine receptors. After embryos reached the tadpole stage (S45), they were fixed with formaldehyde 4% in PBS 1X (v/v) at room temperature during 90 min. Then, washed with PBS 1X three during 5 min. Tadpoles were incubated with glycine 0,15 M in PBS 1X during 30 min. and permeabilizated with PBST (PBS 1X/Triton X-100 1%(v/v)) 10 times during 10 min.

Tadpoles were blocked with PBST-BSA 4% (p/v) overnight at 4°C. Incubation with primary antibody neurofilament 3A10 (Developmental Studies Hybridoma Bank, Iowa, USA) (1:100) in PBST-BSA 4% during 4 days at 4°C. Tadpoles were washed several times with PBST in agitation during 2 h. Then incubated with secondary antibody (1:200) along with αBTX Alexa488 (1:500) in PBST-BSA 4% (p/v) overnight 4°C. Finally, tadpoles were incubated with DAPI (1:100) during 30 min. and were washed several times with PBST During 2 h. and finally washed with PBS 1X. Tadpoles were mounted in glycerol/PBS 1X 50% (v/v).

Z-stack pictures were acquired in a Leica SP8 LIGHTING Spectral confocal microscopy on the CMA Bio-Bio facility, Universidad de Concepción, Chile. Maximal intensity projection images were reconstructed using the Fiji software (Schindelin et al. 2012).

To analyze the axonal branching, we extract the digital skeleton from the Z-stack pictures acquired previously and manually refined and analyzed for the number of branches, junctions, end-point voxels (interpreted as terminal boutons), slab voxels, average branch length, triple, quadruple points and maximum branch length using the AnalyzeSkeleton plugin (Image J, USA). The analyses of overlap between bungarotoxin and axonal fluorescent stain was performed using Fiji version of Image J (Schindelin et al. 2012; Arganda Carreras et al. 2010).

## Results

### Presence of Purinergic receptors during *Xenopus laevis* neurulation

RT-qPCR analysis showed the presence of mRNA for both ionotropic purinergic receptor subunits (X_2,4,5_) and metabotropic purinergic receptor subunits (Y_1,4,6,11,12_) during early (stage 12.5, S12.5), mediate (stage 14, S14) and late (stage 20, S20) neurulation in *Xenopus laevis* (Supplementary figure 1). Relative mRNA expression analysis showed changes in ionotropic receptor subunit expression throughout neurulation. Specifically, P2X2 and P2X5 transcript levels significantly decreased their relative expression to 0.2 and 0.1-fold in S14 and subsequently increased to 10 and 8-fold in S20 respectively, relative to S12.5. In contrast, the P2X4 expression remains largely unchanged between S12-S14 followed by a slight non-significant decrease to 0.5-fold in S20 (Figure 2A). Transcriptional analyses of metabotropic subunits which are insensitive to IVM, showed an increased expression of P2Y11 during S14, whereas the level of expression of P2Y_1,4,6,12_ subunits remained relatively constant. However, during S20 the expression of all subunits, except for P2Y1, statistically increases (Supplementary figure 1). These results show the presence of several purinergic receptors during *Xenopus* neurulation suggesting a relevant contribution of purinergic signaling mediated by ionotropic receptors throughout this developmental process. P2X4 was the only subunit that maintained a relatively stable expression across all neurula stages, possibly associated with a sustained role during neurulation. In addition, P2X4 was strongly immunodetected in whole embryos lysates during early and late neurula stages (S12.5-S20) of *X. laevis*, further supporting its involvement in neurulation (Figure 2B).

**Figure 2.**
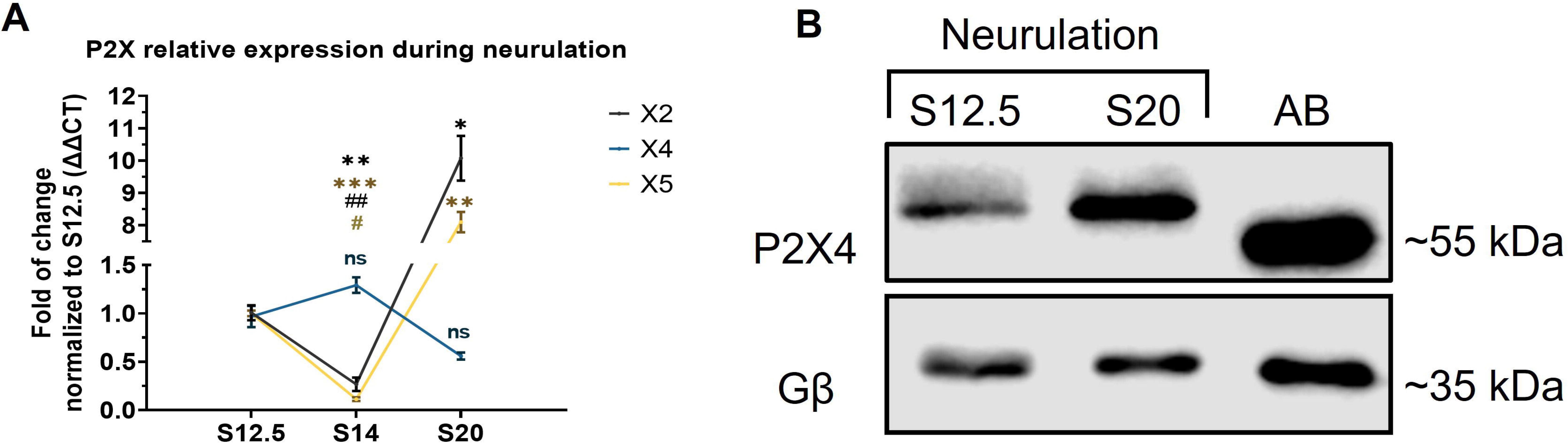
Presence of purinergic subunits in *Xenopus laevis* whole embryo. **(A)** RT-qPCR analysis of relative expression of ionotropic P2X subunits transcripts in neurulation stages. Asterisk (*) indicate comparation between S12.5 and other neurulation stages in the same subunit. # Symbol indicate comparation between X4 subunit and other subunits in S14. N=3. Brown-Forsythe and Welch ANOVA, Dunnet’s correction, (#/* P<0.05, ##/** P<0.01, *** P<0.005). The error bars of the data correspond to the standard deviation of the mean (SEM). **(B)** P2X4 Western blot assay from whole embryo lysates at early (S12.5) and late (S20) neurula with adult brain (AB) as positive control. Gβ shown as loading control N=3.

### Molecular Docking with *Xenopus laevis* P2X4 receptor

To complement the previous results, molecular docking analyses were performed to evaluate interactions between IVM and the transmembrane region of the P2X4 receptor. Two principal docking clusters were identified and are represented in Figure 3A’. Docking simulations using the *X. laevis* P2X4 model revealed two potential interaction pockets (I and II), enumerated according to their binding affinity (kcal/mol) in table 2. The ten highest-scoring conformations were further analyzed using Discovery Studio software. Among these conformations, pocket II predominated, representing 60% of the top-ranked docking poses. Additionally, the average binding affinity of the ten best conformations obtained for *X. laevis* P2X4 model (-8.78 kcal/mol) was compared with that previously reported for the rat P2X4 model (-6.85 kcal/mol), whose interaction had been experimentally validated through electrophysiological analyses (Latapiat et al. 2017).Interaction analysis of the ten highest-scoring conformations revealed that most interacting atoms were located within TM2 (16 of 19). These TM2 atoms were consistently interacting across all conformations and accounted for approximately 75% of the total of interactive atoms identified. Analyses of 2D diagrams excluding van der Waals interactions revealed prevalence of hydrophobic interactions (Alkyl: 10; Pi-alkyl: 6) and carbon-hydrogen bonds (9), with only a single conventional hydrogen bond detected. A key distinction between the two binding pockets was the presence of Pi-alkyl interactions which were absent in pocket I (Figure 3B-B’) but observed in pocket II (Figure 3C-C’). The specific hydrophobic residues from each subunit interacting with IVM across the different conformations are detailed in table 3. Residues identified from the previously published functional P2X4 rat receptor (Latapiat et al. 2017) exhibit high homology with the corresponding region of our generated *X. laevis* model. In TM1, the conserved residues corresponded to Y44, W48 and W52, whereas in TM2 the corresponding residues were M337, S342, G343, and G348 as shown in the sequence alignment shown in Figure 3D.

**Figure 3.**
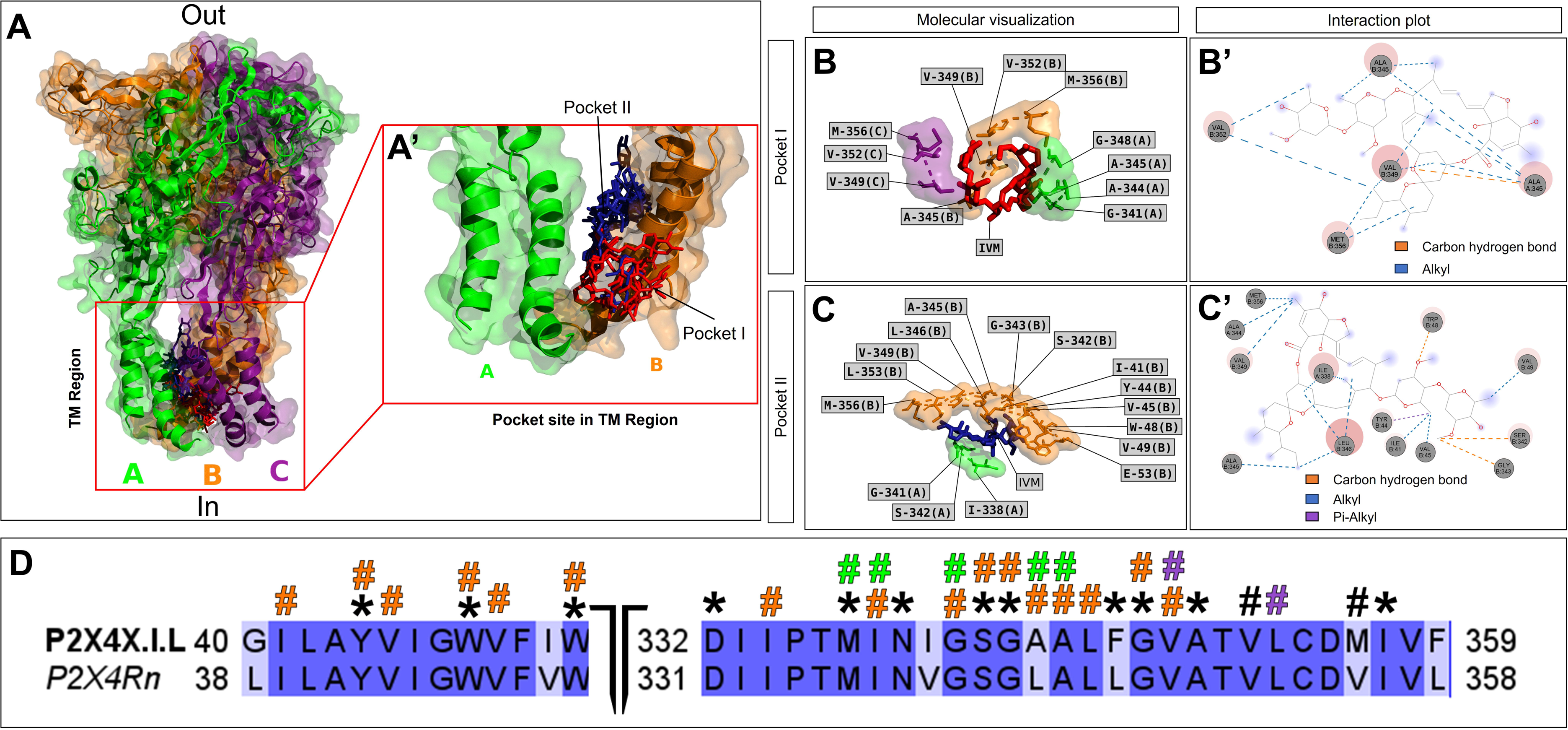
Top scored conformations in transmembrane region of P2X4 Xenopus laevis receptor. **(A)** The receptor is visualized with the extracellular (out) side upwards. Molecular docking of P2X4 structure binding with IVM, presenting the top 10 best scored conformations between P2X4 and IVM which is represented in red for pocket I and blue for II respectively. The three P2X4 subunits A (green), B (orange) and C (purple); are represented in different colors. **(A’)** Structure of TM domain region and conformations formed without 3rd subunit to allow visualization of the interactions. **(B)** Molecular visualization of Subunits forming pocket I with their residues labeled and IVM colored in red. **(B’)** 2D Diagram of interacting residues of pocket I, indicating the type of interaction. **(C)** Molecular visualization pocket II forming Subunits with their residues labeled and IVM colored in blue. **(C’)** 2D Diagram of interacting residues of pocket II, indicating the type of interaction with IVM colored in red. **(D)** Alignment of the transmembrane region of P2X4Xl.L and P2X4Rn, colored according to the percentage of identity. Asterisks (*) show amino acids that appear in validated functional P2X4 models, and (#) symbol indicates residues interacting with IVM. black (#) indicate residues present in the three subunits.

**Table 2.**
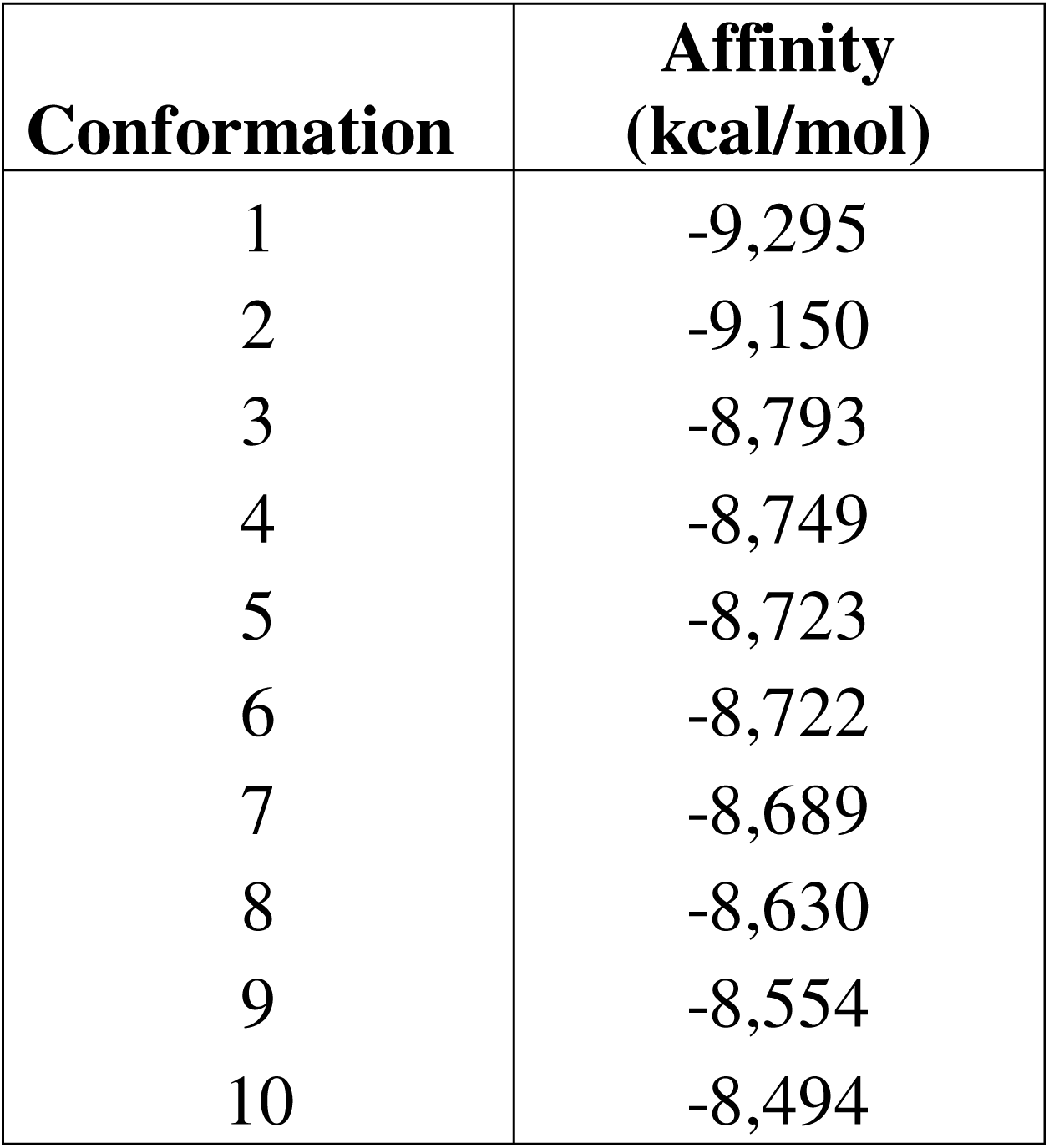
Summary table of best conformations ordered by their affinity.

**Table 3.** Summary table of residues from *Xenopus laevis* model interacting with IVM.

| Residues interacting with IVM |
| --- |
| I41 |
| Y44 |
| V45 |
| W48 |
| V49 |
| W52 |
| M337 |
| I338 |
| G341 |
| A344 |
| A345 |
| L346 |
| G348 |
| V349 |
| V352 |
| L353 |
| M356 |

### Molecular Dynamics Simulations of IVM-mediated Positive Allosteric Modulation of the *Xenopus laevis* P2X4 Receptor

To further characterize the binding of IVM within the previously identified cavities, Molecular Dynamics (MD) simulations were performed using the Desmond (Schrödinger, 2025®). RMSD analysis, utilized to assess the average atomic displacement relative to the initial reference frame, indicated that the protein remained stable, with values fluctuating between 2.0 Å and 2.8 Å (Figure 4A). Regarding the ligand, IVM exhibited oscillations between 2.5 Å and 4.0 Å in Pocket I. In contrast, Pocket II showed a significantly higher displacement, reaching values up to 10.5 Å, suggesting more pronounced conformational rearrangements (Figure 4B). Nevertheless, considering the chemical structure of IVM, both binding pockets exhibited overall stable ligand-receptor conformations during simulations.

**Figure 4.**
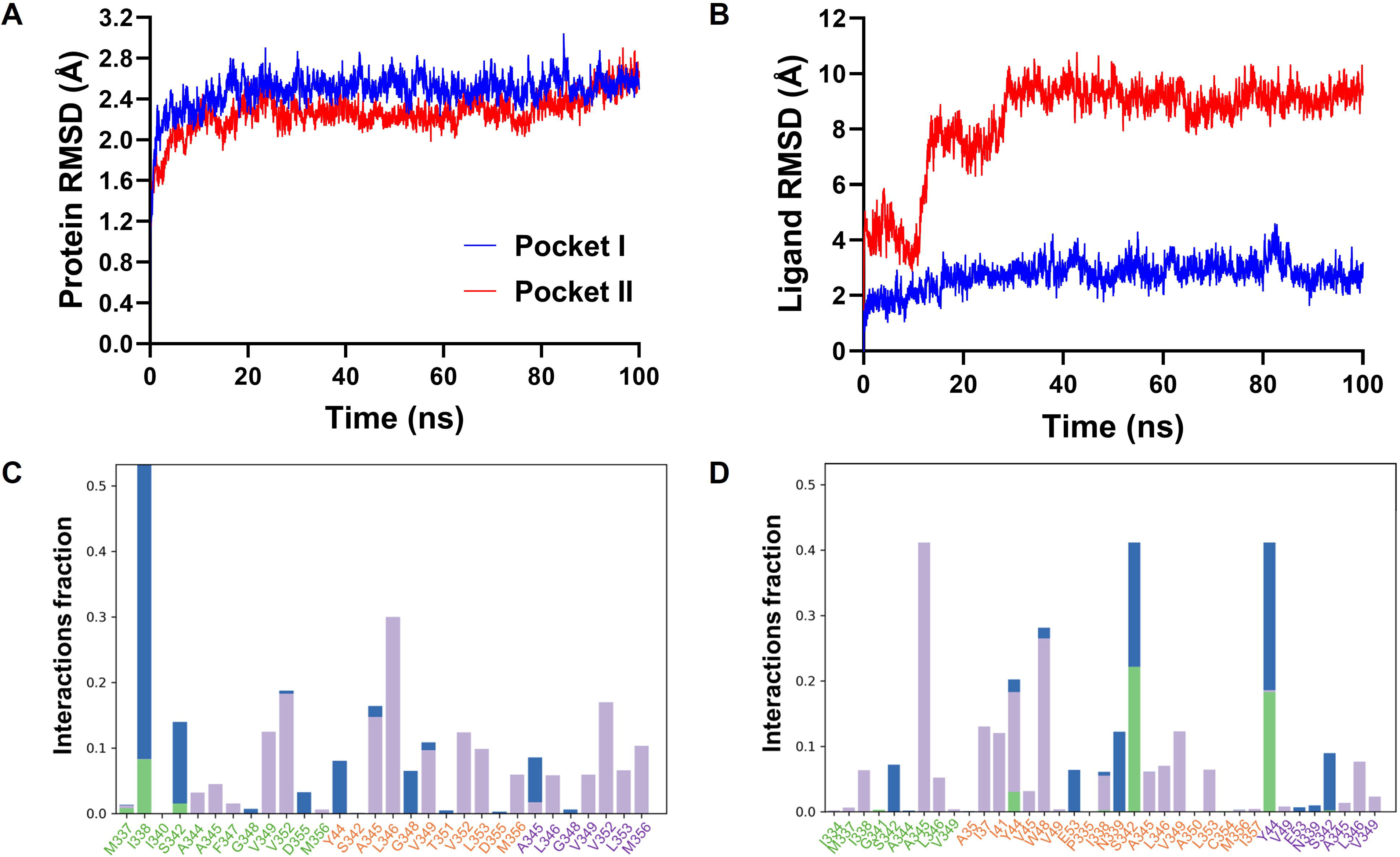
Molecular dynamics simulations for both binding pockets. **(A)** Protein RMSD comparison between both pockets during the simulation. **(B)** RMSD of IVM during the simulation in each binding pocket. **(C)** Protein–ligand contacts analysis for (C) Pocket I and (D) Pocket II, showing interacting residues classified by interaction type and persistence during the simulation. Each bar is colored according to the type of interaction: green indicates hydrogen bonds, blue denotes water-mediated interactions, and purple represents hydrophobic interactions. On the X-axis, residues labels are colored according to receptors subunits: green (subunit A), orange (subunit B) and purple (subunit C).

Residue contact analysis revealed that interactions within Pocket I were predominantly hydrophobic, with residues A345, L346, and V352 contributing to ligand binding during approximately 18% to 30% of the simulation time. Additionally, a water-mediated interaction bridge with I338 was observed for approximately 40% of the trajectory (Figure 4C). Similarly, in Pocket II hydrophobic interactions were also prevalent, specifically involving residues Y44, W48, and A345 which interacted between 20% and 45% of the simulation time. However, hydrogen bonds and water-mediated bridges were also identified involving residues Y44 and S342, which persisted for 20% to 25% of the simulation (Figure 4D). These findings corroborate the molecular docking predictions and highlight the ability of IVM to form structurally and energetically favorable interactions with the receptor.

These results confirm that IVM could bind favorably to *X. laevis* P2X4 receptor, suggesting a potential role as a positive allosteric modulator, similar to previous investigations (Mackay et al. 2017; Illes et al. 2021; Latapiat et al. 2017).

### IVM exposition during neurulation induces tadpole alterations

With the aim of evaluating the effects of P2X4 modulation during neurulation, we examined embryos at the end of neurula stage (S20) to examine possible NTDs and later at tadpole phase (S45) to identify possible secondary developmental consequences. In order to assess the specific participation of P2X4-purinergic receptors, we used IVM a PAM of this specific purinergic channel that significantly increases their Ca^2+^ conductance initiating its modulatory effects from concentrations as low as 250 nM, increasing P2X4 unitary conductance ∼20% (15.3 ± 0.7 pS) and enhancing maximal current by nearly 5-fold (Priel and Silberberg 2004; Mackay et al. 2017). Embryos were subsequently treated with increasing concentrations of IVM (0,1; 0,5; 1; 3; 10; 30; 100 µM) from stage 12.5 until reaching the final neurula stage (S20). IVM exposure did not appear to affect external neural tube closure as no evident morphological differences were observed compared to the control groups (MMR and DMSO) (Figure 5A). However, external phenotypic evaluation at tadpole stages (S45) showed an important increase in pigmentation and paralysis of tadpoles exposed to IVM during neurulation. Analyses of the hyperpigmented phenotype tadpoles showed that the effect exhibited a concentration responsive behavior with a calculated median effective concentration (EC50) of 513 ± 71.7 nM (Figure 5B). Subsequently, we performed a more detailed analysis using two specific concentrations (1 and 10 µM) of IVM (Figure 5C). Treatment with 1 µM IVM increased both the number of melanocytes (95,60 ± 6,877) and the melanocyte-covered area within an arbitrary selected interocular region (49,20 ± 20,80 µm^2^) compared with controls (67.8 ± 2,5 melanocytes and 15,6 ± 3,3 µm^2^). Treatment with 10 µM IVM further increased melanocyte number (118,4 ± 22,20 melanocytes) and covered area (101,4 ± 10,31µm^2^) in comparison with controls, additionally showing the presence of ectopic melanocytes (Figure 5D-E). With the aim to evaluate proper CNS function we assessed two physiological parameters: spontaneous mobility of the tadpoles (S45) and their sensory capacity via escape reflex (Roberts 1990; Spencer et al. 2019). The tadpoles were classified according to their motor behavior in the absence of mechanical stimulation, reflecting basal neuromuscular function (Jamieson and Roberts 2000), and compared with untreated control animals. Control tadpoles exhibited a robust escape reflex in response to stimulation with metal probe. In contrast, tadpoles treated with 1 µM IVM displayed significantly reduced mobility (55,71% ± 13,94) and a subset showed total paralysis (32,85% ± 14,75). The tadpoles exposed to 10 µM IVM predominantly exhibited paralysis and absence of response to stimulation (61,42% ± 16,96), while a smaller proportion displayed reduced mobility (38,57% ± 16,96) (Figure 6). Representative examples of escape reflex responses and locomotor behavior under each experimental condition are shown in Video 1. Due to the paralysis observed in treated tadpoles, we analyzed the internal structure of the CNS in 10 µM IVM neurula-treated tadpoles at stage (S45). We obtained transversal sections of these tadpoles at two different anteroposterior levels and stained them with both the fluorescent dye (DAPI) and hematoxylin-eosin (HE). These results revealed an open neural tube. Notably, this neural tube aperture appears exclusively in the anterior region whereas the posterior region resembled the morphology observed in control animals (Figure 7). These results suggest a concentration-dependent effect of IVM on the CNS of tadpoles. Beginning at 0,5 µM and up to 10 µM we can observe a steadily rise in the number of melanocytes. Consistent with this pattern, a similar concentration-dependent effect was observed in tadpole locomotion. Tadpoles treated with 1 µM IVM exhibited partially reduced mobility, whereas animals exposed to 10 µM displayed a markedly paralyzed phenotype.

**Figure 5.**
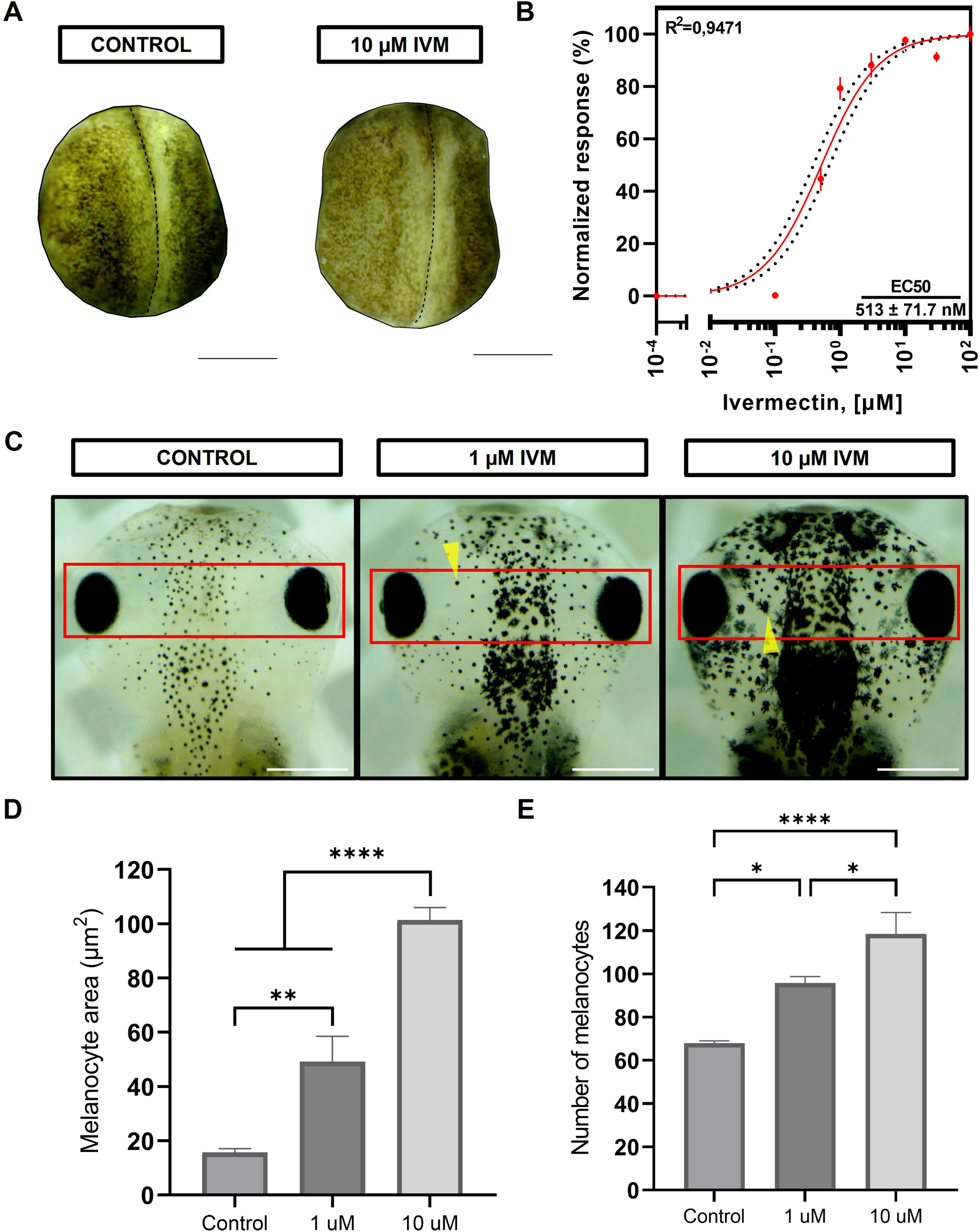
Evaluation of late neurula and tadpole stages exposure to increasing concentrations of IVM. **(A)** Analysis at stage 20 embryos neural tube closure phenotype between DMSO and 10 µM IVM conditions. **(B)** The cover pigmentation produced by melanocytes using increasing concentrations was analyzed from the interocular region (red box) and graphed in a concentration-response curve. **C)** Representative S45 tadpoles shown an increase of pigmentation in response to different concentrations of IVM. Ectopic melanocytes are marked with yellow arrows. **(D)** Graph shows the analysis of melanocyte area in the analyzed conditions. **(E)** Graph shows the analysis of number of melanocytes in the analyzed conditions. N= 5 One-way ANOVA Dunnett correction (* P<0.05, **P<0.01, **** P<0,0001). The error bars of the data correspond to the standard error of the mean (SEM). Scale bar: 0.5 mm.

**Figure 6.**
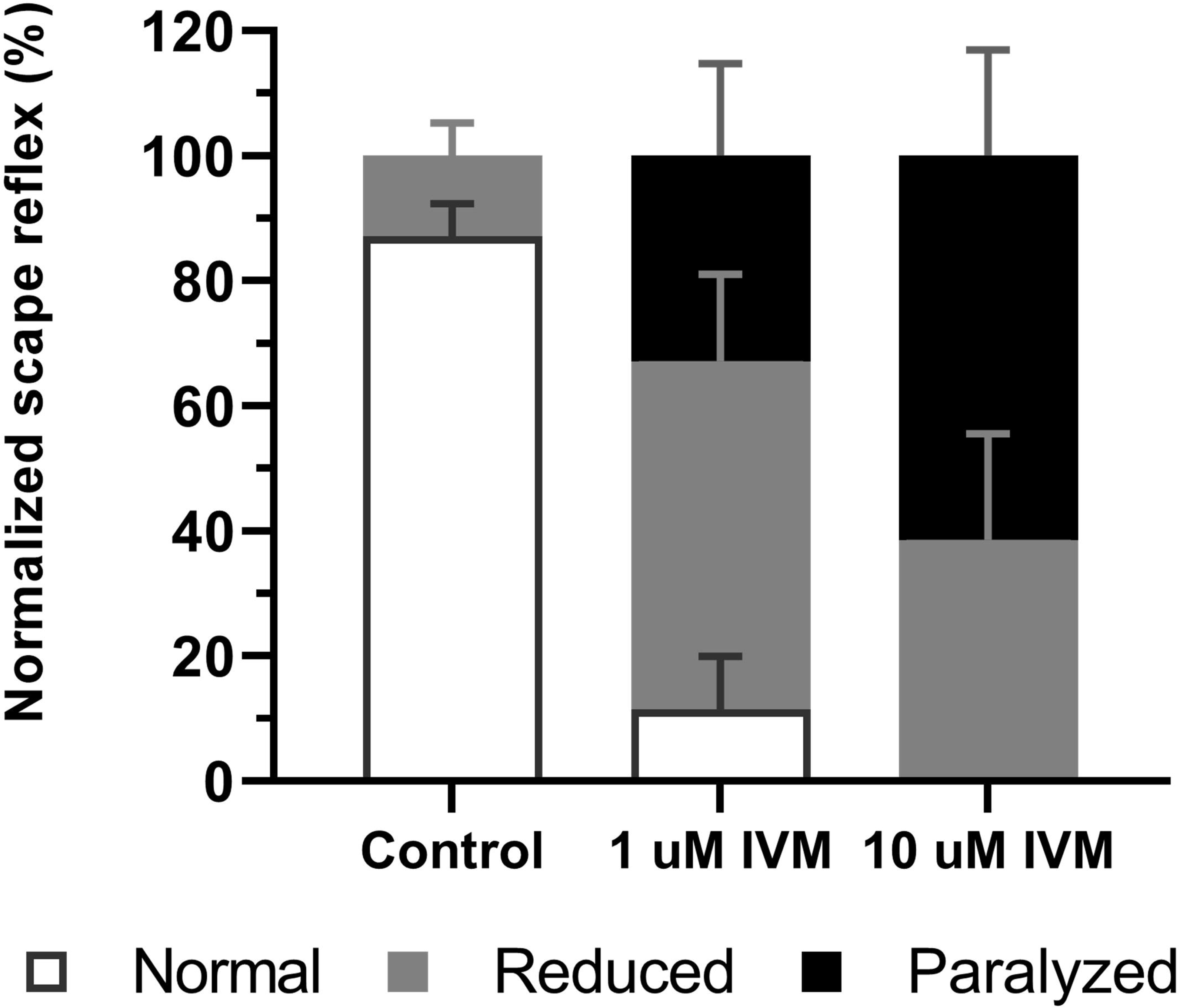
Effect of IVM on touch-evoked escape reflex of *Xenopus laevis* stage 45 tadpole. Tadpoles were assigned to three groups depending on their response: Normal (white), reduced (grey) and paralyzed (black). Results are reported as percentage of total batches of embryos analyzed. Number of batches analyzed = 7, 10 embryos per condition. The error bars of the data correspond to the standard error of the mean (SEM). (Spencer et al. 2019)

**Figure 7.**
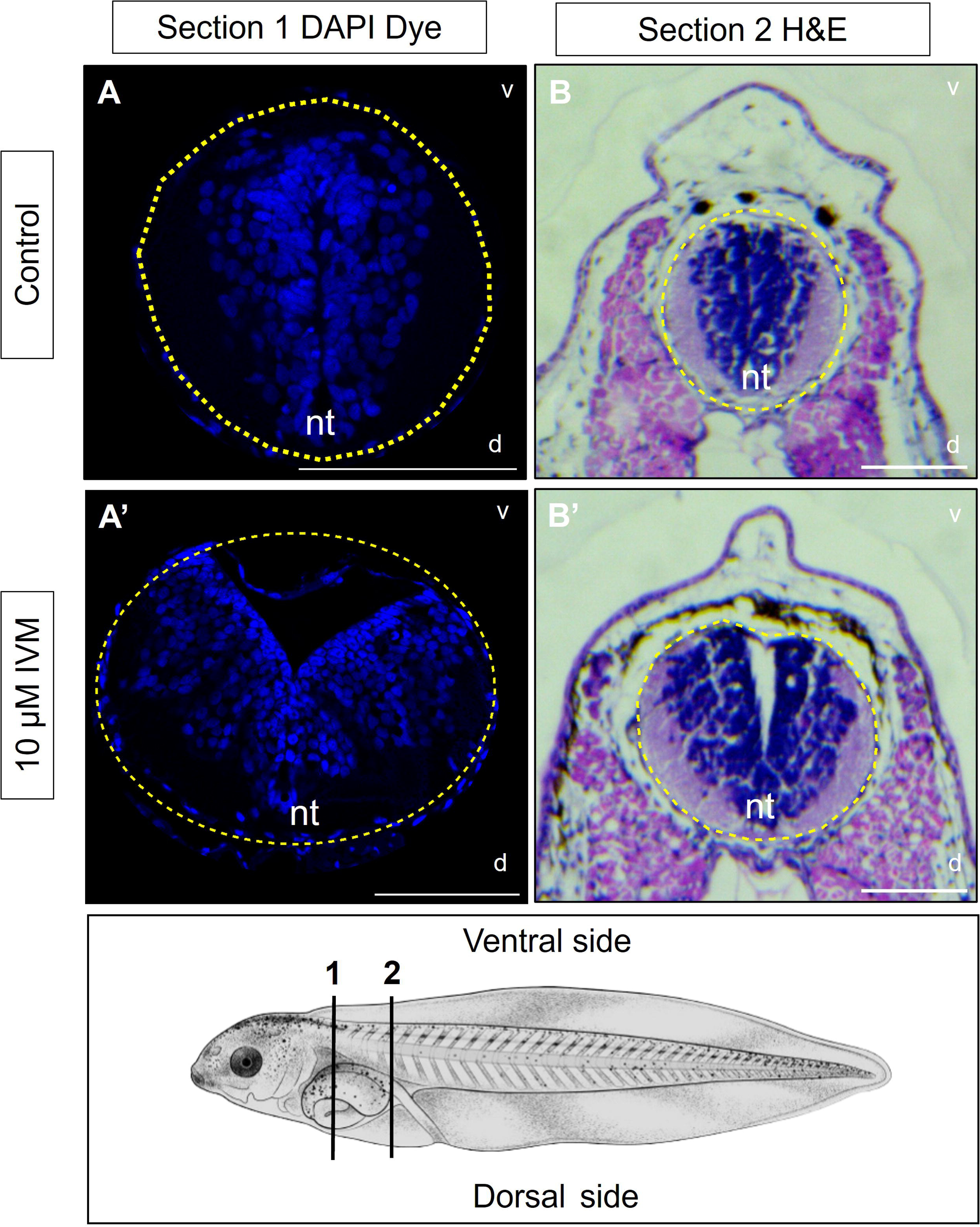
Early exposure to IVM causes open neural tube defects. Histologic techniques on different *X. laevis* tadpole S45 transversal sections. Section 1 were processed with immunofluorescence DAPI. Section 2 were processed with hematoxylin-eosin. **(A-B)** Sections on control tadpoles., **(A’-B’)** The IVM effects are observed in the 10 µM IVM. d= Dorsal, v= Ventral, nt= Neural tube. Scale bar = 100 µm.

To further investigate the paralysis phenotype, we evaluated the expression of three major molecular markers associated with motoneuron (MN) differentiation by qPCR. Specifically, we analyzed mRNA expression levels of transcription factors involved in MN formation and development, including MNX1, a transcription factor required for MN differentiation; NCAM, a molecule involved in axonal guidance and targeting that is regulated through transcriptional modification; and NKX6.1, required for proper MN axonal growth, whose morpholino-mediated knockdown has previously been associated with paralysis phenotypes in embryos (Arber et al. 1999; Dichmann and Harland 2011; Hinsby 2004). However, no statistically significant differences were observed between control tadpoles or those treated either 1 or 10 µM IVM (Supplementary figure 2).

### Evaluation of additional cys-loop receptors target of IVM

Several pharmacological studies have reported that IVM can modulate receptors, such as GlyR, GABA(A)R, and nAChR. Because GlyR and GABA(A)R mediate inhibitory signaling, we evaluated their expression before neurulation (S10), during neurulation (S12.5, S14, and S19) and in adult brain (AB) tissue. In addition, we performed PCR analysis of VIAAT (Vesicular Inhibitory Amino Acid Transporter), a synaptic vesicle protein responsible for the vesicular accumulation of inhibitory neurotransmitters such as GABA and glycine. We also evaluated the expression of target receptor subunits including GlyR-α2, GABA(A) α1 and α2. Our results showed an absence of VIAAT expression during neurulation, whereas expression was clearly detected in adult brain samples (Figure 8A), suggesting the absence of functional GABAergic and glycinergic signaling during early stages of *Xenopus* development (Saito et al. 2010). Nevertheless, the GlyR-α2 subunit was detected both during neurulation and in adult brain tissue. To complement the analysis of inhibitory signaling during neurulation, we evaluated the role of GlyR using strychnine (Stry), a competitive GlyR antagonist that binds to the orthosteric site of the receptor (Zlotos et al. 2022). Because GlyR has previously been proposed as a potential mediator of IVM effects in *Xenopus*, we extended our analysis to later developmental stages. Under these conditions, no evident morphological alterations were observed at any of the developmental stages evaluated. Embryos exhibited normal neural tube closure and typical post-neurula development. Furthermore, analyses of body length, mobility and pigmentation in stage 45 (S45) tadpoles revealed no significant differences compared with control animals (Figure 8B-F). Analyses of GABAergic signaling, consistent with previous data reported, did not detect expression of GABA(A) α1 subunit during neurulation stages, whereas the α2 subunit was detected (Kaeser et al. 2011) (Figure 8A). Then we evaluated a potential role of GABA(A)R during the neurulation process by performing treatments with different concentrations of bicuculline, a specific channel blocker and GABA(A)R antagonist (Johnston 2013). We did not observe any morphological alterations in neural tube closure at stage 20, neither were morphological abnormalities nor differences in tadpole length detected at stage 31 (Figure 8G-I).

**Figure 8.**
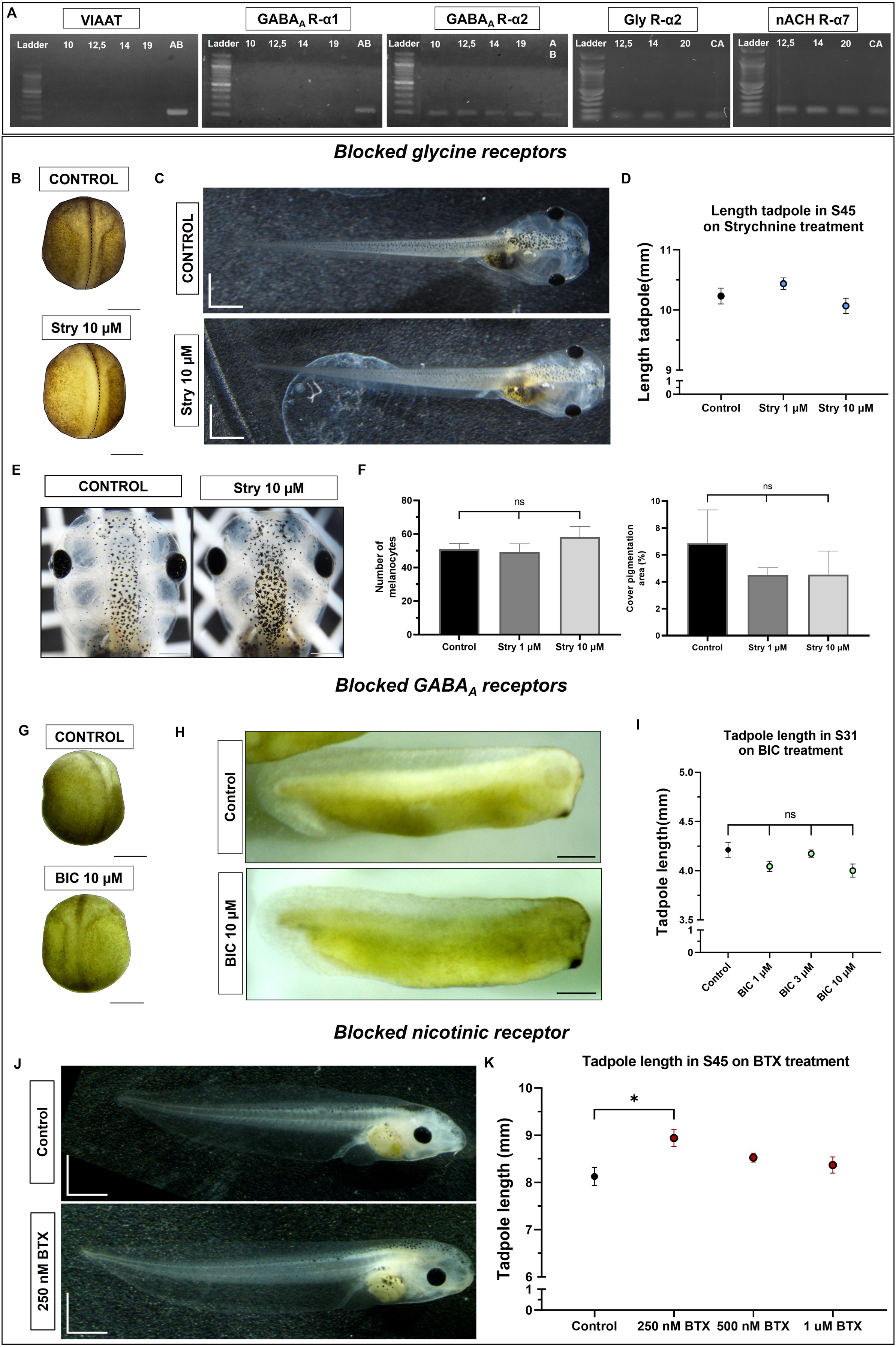
Evaluation of ionotropic receptors IVM-modulated. **(A)** PCR of VIAAT (Vesicular Inhibitory Amino Acid Transporter), GABA(A) α1, GABA(A) α2, Gly R α2 and nACh R α7 subunit on neurulation stages. Products were separated by agarose gel electrophoresis. Ladder = 100 bp **(B)** Analysis at stage 20 embryos neural tube closure phenotype between control and 10 strychnine µM. **(C)** Picture at S45 between control and Stry phenotype. **(D)** Plotted values of tadpole length in S45 on Stry treatments, N=5. **(E)** Interocular dorsal pigmentation comparison S45 between control and Stry treatment. **(F)**. Plotted interocular pigmentation analysis on number of melanocytes (left panel) and cover pigmentation (right panel), N=5. **(G)** Picture at stage 20 embryos neural tube closure phenotype between control and 10 bicuculline µM. **(H)** Picture at S45 between control and BIC phenotype. **(I)** Plotted values of tadpole length in S31 on BIC treatments, N=5. **(J)** Picture at S45 between control and BTX phenotype. **(K)** Plotted values of tadpole length in S45 on BTX treatments, N = 9, (* P<0.05), D, I, K: Brown-Forsythe and Welch ANOVA test. F: Ordinary one-way ANOVA test. B, E, G and H scale bar: 0.5 mm. C, J scale bar: 1mm. The error bars of the data correspond to the standard error of the mean (SEM).

Lastly, we performed complementary experiments to evaluate the presence and potential involvement of cholinergic excitatory signaling during neurulation through nicotinic α7 receptors. PCR analyses revealed the presence of the α7 subunit throughout all neurulation stages as well as in adult brain tissue (Figure 8A). To investigate its role during neurulation, embryos were treated with bungarotoxin (α-BTX), a potent and irreversible competitive antagonist with partial selectivity for nicotinic α7 receptors (Pohanka 2012), at concentrations of 250 nM, 500nM and 1µM. Tadpoles treated with 250 nM α-BTX exhibited a significant increase in body length compared with control animals (Figure 8J-K). In addition, a low incidence (∼10%) of developmental abnormalities was observed in embryos treated with 250 nM and 500 nM α-BTX, including generalized edema, alterations in the muscle chevron pattern, and an appendix projection. However, these abnormalities were not observed in embryos treated with 1µM α-BTX. Finally, and similar to previous evaluations, we did not find changes in pigmentation, tube closure, or mobility. Taken together, these findings indicate that none of the alternative signaling pathways evaluated reproduced the phenotypes induced by IVM treatment, supporting a predominant role for purinergic P2X4 signaling during neurulation.

### Analyses of neuromuscular structure in S45 Tadpoles using Whole-mount fluorescence immunohistochemistry

Considering the previous analyses of pigmentation, neural structure, transcription factor expression, and particularly locomotor behavior, we further evaluated the neuromuscular junction (NMJ) structure in detail following treatment of embryos with 10 µM IVM, the condition that produced the strongest motor phenotype.

To identify and analyze NMJs in tadpoles, whole-mount immunofluorescence staining was performed and compared with control embryos. The 3A10 antibody (1:100) was used to label neurofilament-associated proteins in motor neurons, allowing clear visualization of axonal tracks in S45 *X. laevis* tadpoles (Bermedo-García et al. 2018) (Figure 9A). In parallel, we used alpha-bungarotoxin (α-BTX) to label the AChRs, giving a strong postsynaptic staining. Initially, both control and treated tadpoles exhibited a comparable innervation pattern. However, detailed skeletonized analyses revealed that IVM-treated tadpoles displayed a significant increase in axonal branching (Figure 9B’’-C’’). This effect was consistently observed in six out of nine parameters associated with presynaptic NMJ organization. In particular, treated tadpoles exhibited increased numbers of branches (1.93 ± 0.24 vs 1.12), junctions (2.02 ± 0.27 vs 1.00), and terminal boutons (2.34 ± 0.26 vs 1.00) without significant changes in average branch length (1.08 ± 0.14 vs 0.94). These results suggest that axons in treated tadpoles generate a greater number of new branches rather than elongating (consolidating) pre-existing ones. Additionally, the increase in the number of terminal boutons pixels is showing that the NMJ is more complex and arborized than the control. In contrast, no significant differences were observed in the expression or distribution of postsynaptic staining, resulting in a significant reduction of synaptic contact, measured as overlap between neurofilament/bungarotoxin (0.7 ± 0.06 vs 0.93) (Figure 9D).

**Figure 9.**
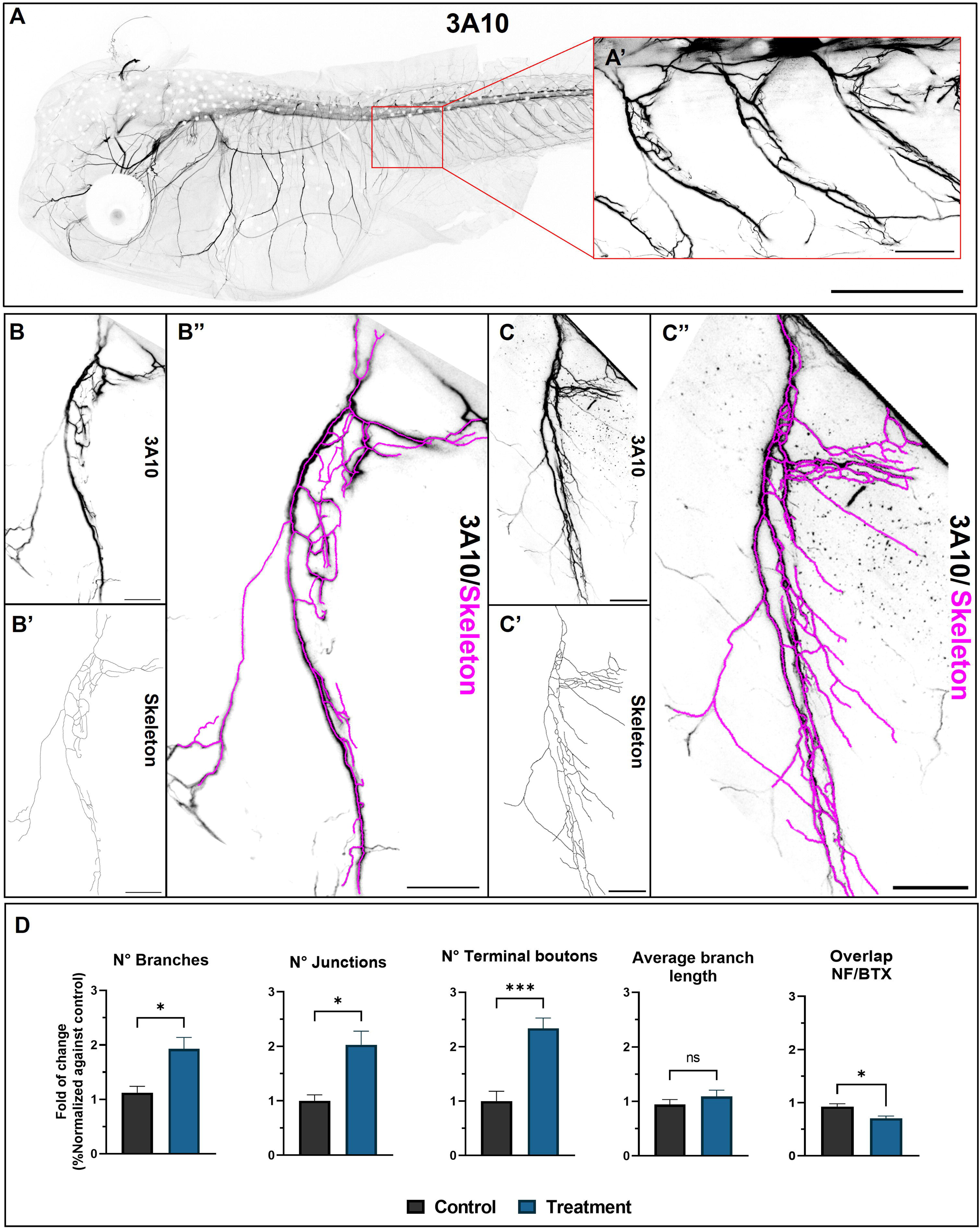
Immunolabeling of neurofilaments (3A10) in Whole mounted *X. laevis* at the trunk ventral region in tadpoles (Stg 45) imaged by confocal microscopy. (A) S45 control tadpole stained for the neurofilament-associated specific marker 3A10. **(A’)** 40X Zoom at trunk ventral region. **(B)** Innervation in control and treatment (IVM 10 µM) **(C)** tadpoles of neurofilaments. Illustration of axonal branching thought skeleton **(A’-B’).** Merge of skeleton and innervation in control **(B’’)** and treatment **(C’’)** tadpoles. **(D)** Number of branches, junction, terminal boutons and average branch length obtained via AnalyzeSkeletonize plugin and overlap of neurofilament v/s bungarotoxin in control and neurula-treated tadpoles. N=5. Welch’s t test (* P<0.05, *** P<0.001). The error bars of the data correspond to the standard error of the mean (SEM). A Scale bar: 1mm. A’, B-B’’, C-C’’ Scale bar: 100 µm.

## Discussion

Previous studies have characterized the expression of purinoreceptors during embryogenesis, suggesting morphogenic roles beyond their well-established pharmacological and structural functions, particularly for P2X and P2Y receptor subtypes (Burnstock and Ulrich 2011; Ulrich and Illes 2014). Moreover, dysfunctional ionotropic P2X receptors have been identified with several pathological conditions, including hearing loss, neuropathic pain, and inflammatory bone loss, highlighting their clinical relevance (Illes et al. 2021).

Purines and pyrimidines are recognized as important signaling molecules during embryonic development across a wide range of biological models, including humans, particularly in the development of the nervous system. However, unlike metabotropic receptors, the specific roles and mechanism of most ionotropic purinergic receptors during processes such as neural tube formation remain poorly understood.

In the present study, we analyzed the dynamic mRNA expression profiles of P2X_2,4,5_ receptors throughout neurulation. Our results revealed a sustained expression of the P2X4 receptor, the purinergic channel with the highest calcium conductance within the P2X family, whereas P2X_2,5_ transcript levels decreased during the intermediate neurula stage. In addition, western blot analyses demonstrated strong P2X4 protein expression in whole embryos.

These results suggest an important role for P2X4 throughout the neurulation process, potentially associated with its role in Ca^+2^ mobilization during early vertebrate development. Calcium signaling has been implicated in several key developmental processes in *X. laevis* embryos, including cell proliferation, migration, and apical constriction, which will be further investigated in future studies (Goyal et al. 2026; Moreau et al. 2016; Sequerra et al. 2018).

### IVM concentration-response on hyperpigmentation

With the objective of modulating P2X4 receptor activity, and based on previous studies, we used IVM during early embryonic development, specifically throughout the neurulation period. This experimental approach allowed us to determine pharmacological parameters such as EC50 and establish a concentration-response effect between IVM exposure and tadpole hyperpigmentation, consistent with previous reports.

This phenotype was not detected at 100 nM IVM but became evident at concentrations beginning at 500 nM. This observation suggests that the effects of IVM above the 250 nM threshold are consistent with the reported pharmacological sensitivity of P2X4 receptor modulation (Priel and Silberberg 2004).

We postulate that hyperpigmentation effect observed in this study is unlikely to be mediated by the glycine receptor chloride channel (GlyCl), as proposed by Blackinston et al. If the effects of IVM were primarily GlyCl-dependent, a pigmentation response would be expected at concentrations around 100 nM or less, given that GlyCl and also GABA_A_ receptors potentiation by IVM is known to initiate at nanomolar concentrations; 30 nM (EC_50_=0.39 µM) and 1 nM (EC_50_=17.8 nM) respectively (Shan et al. 2001; Krůšek and Zemková 1994).

In contrast, the threshold to increase α7 current begins at micromolar concentrations of IVM (3 µM, EC50=6.8±1.2 µM), making its participation unlikely at the lower concentrations used in this investigation (Collins and Millar 2010). Therefore, the absence of detectable effects at 100 nM argues against a predominant role for GlyCl and GABA(A) receptors under our experimental conditions, although it does not completely rule out their possible contribution. Instead, our results raise the possibility of another IVM target similar to P2X4, which is modulated at similar concentrations to our observed concentration-response effects.

### IVM lead to NMJ alterations and spinal cord analysis

Additionally, to the pigmentation response, Blackiston et al., also described a paralyzing effect on IVM-treated embryos in more prolonged treatments spanning stages 13-30. Our results show that the use of IVM exclusively during neurulation (S12-20) also causes motor paralysis on later tadpole stages (S45) and, similar to pigmentation, this effect seems to be concentration dependent as tadpoles treated with 1 µM IVM during neurulation exhibited reduced escape reflex responses and, in a smaller proportion of animals, complete paralysis. This phenotype became more pronounced following treatment with 10 µM IVM (Figure 6).

Due to the severity of the observed motor phenotype, characterized by pronounced paralysis, we performed an exhaustive whole-mount immunofluorescence analysis of the NMJ in paralyzed tadpoles that had been treated with 10 µM IVM. Our results revealed that IVM exposure induces a hyperinnervation phenotype. This phenotype is characterized by a significant increase in axonal branching compared with control animals, which could disrupt normal synaptic transmission by promoting asynchronous neurotransmitter release, reducing signaling efficiency, altering NMJ stability, and impairing complete synaptic maturation (Huang et al. 2022). The axonal branching dynamics observed in our study resemble those previously described in *D. melanogaster* where dysregulation of EGFR signaling produces excessive branching due to defects in axonal pruning, without major alterations in the motor end plate but accompanied by reduced NMJ overlap (Dutta et al. 2023; Zschätzsch et al. 2014).

Moreover, previous studies have also reported that IVM treatment induces hyperinnervation phenotypes (Blackiston et al. 2015; Cairns et al. 2018). In one of these studies, hyperinnervation was detected only in host embryos were exposed to IVM from the tailbud stage (S24) to the tadpole stage (S46). In contrast, a recently published study reported a hypo-innervation phenotype accompanied by transient paralysis at S45 following exposure of embryos to a lower concentration of IVM (1 µM) over an extended developmental window (S9-S26) (Jones et al. 2025). Taken together, the differences between our findings and previous reports highlight that the mechanisms underlying IVM-induced axonal branching alterations during development remain incompletely understood. Here, we highlight the relevance of the pharmacological exposure window, identifying neurulation as a critical developmental period influencing the establishment of physiological NMJ organization in the chordate *X. laevis*, potentially generating a pharmacological imprinting phenomenon.

Due to the paralysis observed in tadpoles treated with 10 µM IVM, we further evaluated spinal cord organization through transversal sections. Our results revealed an aperture in the anterior region of the spinal cord. This region corresponds primarily to sensory neuronal domains, whereas the ventral region contains motor neuron populations, with the aperture appearing more pronounced in the anterior region of the spinal cord. The apparently normal external closure of the neural tube suggests two options: 1) During primary neurulation (S12-20) one or more mechanical processes involved in neural tube formation, such as apical protrusion, luminal intercalation, or neural fold fusion, may be affected by altered Ca^+2^ signaling (Vijayraghavan and Davidson 2017), or 2) Dysregulation of the neuromesodermal progenitor cell expression during the second phase of neurulation could preferentially impair anterior neural tube organization (Shaker et al. 2021).

### Presence of another IVM target during *Xenopus* neurulation

Despite IVM being considered a selective PAM of the P2X4 receptor, it is also capable of modulating several ligand-gated ion channels and receptors in vertebrates, including nicotinic, GABAergic and glycinergic receptors. Based on our results, we associate the major contribution of the observed phenotypes primarily with P2X4 signaling, and to a lesser extent with GABAergic and glycinergic transmission, due to the absence of VIAAT, a key protein required for effective inhibitory neurotransmission. Even with the presence of inhibitory receptors there is a lack of mechanisms to explain such modulation. Furthermore, no significant phenotypic alterations were detected following treatment with the GlyR and GABA(A) receptor antagonists strychnine and bicuculline during neurulation, further supporting a predominant role for P2X4 signaling in the developmental effects induced by IVM.

Additionally, GABA(A)R modulated by IVM commonly exhibit a pentameric composition α*2*β*2*γ (Estrada-Mondragon and Lynch 2015). Although the IVM modulation properties described for the α1 subunit are also conserved in the α2 subunit, our bibliographical analyses did not detect the presence of the β or γ subunits required for the assembly of a functional GABA(A)R channel during *Xenopus* neurulation.

Another modulable IVM target are the nicotinic acetylcholine receptors (homomeric α7), whose expression together with choline acetyltransferase (ChAT) has been reported during E13 mouse spinal cord development (Phelps et al. 1990). Our results show α7 subunit mRNA transcription during *Xenopus* neurulation, however, blockade of nicotinic receptors did not reproduce IVM-associated phenotypes such as pigmentation changes or locomotor alterations, suggesting that the nicotinic signaling contributes only minimally during neurulation and supporting the involvement of alternative pathways.

Although we do not exclude the potential involvement of other ligand-gated ion channels during neurulation, our findings strongly point toward ionotropic purinergic channels, particularly P2X4 receptors, as major mediators of the effects induced by IVM through Ca^2+^ dependent mechanisms. In addition, our group detected the expression of both ionotropic and metabotropic purinergic transcripts, as well as the vesicular nucleotide transporter (VNUT), further supporting the existence of an active purinergic signaling system during neurulation. This contrasts with the alternative signaling pathways evaluated in the present study. This observation becomes especially relevant when considering the wide range of phenotypes identified both in our work and in previous studies using *Xenopus laevis,* including alterations in muscle patterning after prolonged exposure (Lobikin et al. 2015), vacuolated musculature, digestive system abnormalities in tadpoles (Supplementary Figure 3-4), altered presynaptic NMJ distribution, hyperpigmentation, and craniofacial defects (Jones et al. 2025). Together, these findings suggest a broad developmental impact across multiple systems, likely resulting from a pharmacological imprinting phenomenon occurring during neurulation.

### In silico interaction of IVM on *Xenopus* P2X4 receptor

Our in silico analyses validate our *Xenopus laevis* P2X4 receptor model and demonstrate a favorable binding affinity of IVM for this model. In addition, the analyses revealed a pattern of interacting residues in TM1 and TM2 homologous to those previously described by Latapiat et al. 2017 which were functionally validated using electrophysiology. Furthermore, this study by Latapiat and colleagues evaluated the molecular basis of IVM selectivity, identifying the residues Y42, W46 and W50 on TM1, together with M336, S341, G342 and G347 in TM2 as critical determinants distinguishing P2X4 from other P2X receptor subtypes and generating a favorable environment for IVM modulation. Notably, these residues, as well as the interaction patterns involving pi-alkyl interactions with Y44 in TM1 and hydrogen bonding with S342 in TM2, closely matched the molecular docking results obtained in our study.

The combined data from molecular docking and MD simulations reveal that the key residues involved in the interaction within each binding pocket largely overlap, validating the proposed model system. Moreover, the pronounced ligand RMSD fluctuations observed in Pocket II (reaching up to 10.5 Å) indicate increased IVM mobility within this site, likely associated to the specific amino acid composition and the chemical nature of the residues involved.

The prevalence of hydrophobic interactions is consistent with the lipophilic nature of IVM. Nonetheless, the presence of hydrogen bonds and water-mediated bridges involving Y44 and S342 in Pocket II can be explained by the biochemical properties of these residues. Collectively, these results suggest that Pocket I constitutes a more energetically favorable binding environment for IVM, primarily stabilized through hydrophobic interactions, making it a more likely candidate for stable binding compared to Pocket II.

Additionally, it is possible that IVM may bind to and potentiate heteromeric P2X4/6 receptors. However, the presence of P2X6 subunits during neurulation has not been reported in X*enopus* (Blanchard et al. 2019; Khakh et al. 1999).

### Limitations and future approaches

The P2X4 transcript was reported from the blastula stage in *Xenopus* which compromises the use of more specific molecular approaches such as morpholino-mediated knockdown to evaluate the role of P2X4 during early development. Although we detected the endogenous presence of the P2X4 receptor, its precise location within the embryo during different stages of neurulation, as well as the specific germinal layers or structures affected by IVM modulation, remains unknown. These limitations highlight the need for future studies aimed at elucidating both the spatial distribution of P2X4 receptors and the molecular mechanisms underlying their function during neurulation. Such approaches could be complemented by the use of novel allosteric modulators such as 5-BDBD, parotoxine, BX-430 and BAY1797, among others (Werner et al. 2019). Given the challenge associated with the use of molecular tools, the identification and application of specific pharmacological compounds become especially relevant for investigating P2X4 receptor function, particularly when combined with complementary approaches such as molecular dynamics analyses to further characterize purinergic signaling during the neurulation process.

## Conflict of interest

The authors declare that the research was conducted in the absence of any commercial or financial relationships that could be construed as a potential conflict of interest.

## Author contribution

**C.C.-V:** Writing – original draft, review and editing, Formal Analysis, Methodology, Software, Data acquisition, Data curation, Visualization, Conceptualization, Investigation. **P.A.C:** Writing – original draft, Writing – review and editing, Resources, Project administration, Visualization, Conceptualization, Supervision, Investigation, Validation, Funding acquisition. **C.F.B:** Writing – review and editing, Methodology, Resources, Software. **B.S.-M:** Writing – review and editing, Methodology, Software. **C.V:** Writing – review and editing., Methodology. **N.A.F:** Writing – review and editing, Methodology **G.E.Y:** Writing – review and editing, Resources, **G.M.-C:** Writing – review and editing, Resources.

## Funding

This research was funded by ANID-FONDECYT 1231038 (to P.A.C.), ANID-FONDECYT 1250856 (to G.E.Y.), as well as ANID-FONDECYT 1251488 (to G.M.-C).N.F and B.S-M were supported by the Universidad de Concepción through Graduate School Fellowships (Programa de Magister en Neurobiologia).

## Notes

### Competing Interest Statement

The authors have declared no competing interest.

## References

Andrejew, Roberta, Natalia Turrini, Qing Ye, Yong Tang, Peter Illes, and Henning Ulrich. 2023. “Purinergic Signaling in Neurogenesis and Neural Fate Determination: Current Knowledge and Future Challenges.” In Purinergic Signaling in Neurodevelopment, Neuroinflammation and Neurodegeneration, edited by Henning Ulrich, Peter Illes, and Talita Glaser. Springer International Publishing. 10.1007/978-3-031-26945-5_5.

Arber, Silvia, Barbara Han, Monica Mendelsohn, Michael Smith, Thomas M. Jessell, and Shanthini Sockanathan. 1999. “Requirement for the Homeobox Gene Hb9 in the Consolidation of Motor Neuron Identity.” Neuron 23 (4): 659–74. 10.1016/S0896-6273(01)80026-X.

Arganda□Carreras, Ignacio, Rodrigo Fernández□González, Arrate Muñoz□Barrutia, and Carlos Ortiz□De□Solorzano. 2010. “3D Reconstruction of Histological Sections: Application to Mammary Gland Tissue.” Microscopy Research and Technique 73 (11): 1019–29. 10.1002/jemt.20829.

Bates, Nicola. 2020. “Poisons Affecting the Neurological System.” The Veterinary Nurse 11 (3): 116–25. 10.12968/vetn.2020.11.3.116.

Benavides-Rivas, Camila, Lina Mariana Tovar, Nicolás Zúñiga, et al. 2020. “Altered Glutaminase 1 Activity During Neurulation and Its Potential Implications in Neural Tube Defects.” Frontiers in Pharmacology 11 (June): 900. 10.3389/fphar.2020.00900.

Bermedo-García, Francisca, Jorge Ojeda, Emilio E. Méndez-Olivos, Sylvain Marcellini, Juan Larraín, and Juan Pablo Henríquez. 2018. “The Neuromuscular Junction of Xenopus Tadpoles: Revisiting a Classical Model of Early Synaptogenesis and Regeneration.” Mechanisms of Development 154 (December): 91–97. 10.1016/j.mod.2018.05.008.

Blackiston, Douglas J., George M. Anderson, Nikita Rahman, Clara Bieck, and Michael Levin. 2015. “A Novel Method for Inducing Nerve Growth via Modulation of Host Resting Potential: Gap Junction-Mediated and Serotonergic Signaling Mechanisms.” Neurotherapeutics 12 (1): 170–84. 10.1007/s13311-014-0317-7.

Blanchard, Camille, Eric Boué-Grabot, and Karine Massé. 2019. “Comparative Embryonic Spatio-Temporal Expression Profile Map of the Xenopus P2X Receptor Family.” Frontiers in Cellular Neuroscience 13 (July): 340. 10.3389/fncel.2019.00340.

Boothby, K. M., and A. Roberts. 1995. “Effects of Site of Tactile Stimulation on the Escape Swimming Responses of Hatchling *Xenopus Laevis* Embryos.” Journal of Zoology 235 (1): 113–25. 10.1111/j.1469-7998.1995.tb05132.x.

Burnstock, Geoffrey, and Henning Ulrich. 2011. “Purinergic Signaling in Embryonic and Stem Cell Development.” Cellular and Molecular Life Sciences: CMLS 68 (8): 1369–94. 10.1007/s00018-010-0614-1.

Cairns, Dana M., Jodie E. Giordano, Sylvia Conte, Michael Levin, and David L. Kaplan. 2018. “Ivermectin Promotes Peripheral Nerve Regeneration during Wound Healing.” ACS Omega 3 (10): 12392–402. 10.1021/acsomega.8b01451.

Close, Bryony, Keith Banister, Vera Baumans, et al. 1997. “Recommendations for Euthanasia of Experimental Animals: Part 2.” Laboratory Animals 31 (1): 1–32. 10.1258/002367797780600297.

Collins, Toby, and Neil S. Millar. 2010. “Nicotinic Acetylcholine Receptor Transmembrane Mutations Convert Ivermectin from a Positive to a Negative Allosteric Modulator.” Molecular Pharmacology 78 (2): 198–204. 10.1124/mol.110.064295.

CRUMP, Andy, and Satoshi ŌMURA. 2011. “Ivermectin, ‘Wonder Drug’ from Japan: The Human Use Perspective.” Proceedings of the Japan Academy. Series B, Physical and Biological Sciences 87 (2): 13–28. 10.2183/pjab.87.13.

Dassault Systèmes. 2025. BIOVIA Discovery Studio 2025. V. v.25.1.0.24284. Released.

Delarasse, Cécile, Pauline Gonnord, Micaela Galante, et al. 2009. “Neural Progenitor Cell Death Is Induced by Extracellular ATP via Ligation of P2X7 Receptor.” Journal of Neurochemistry 109 (3): 846–57. 10.1111/j.1471-4159.2009.06008.x.

Diazgranados-Sanchez, J. A., J. L. Mejia-Fernandez, L. S. Chan-Guevara, M. H. Valencia-Artunduaga, and J. L. Costa. 2017. “[Ivermectin as an adjunct in the treatment of refractory epilepsy].” Revista De Neurologia 65 (7): 303–10.

Dichmann, Darwin S., and Richard M. Harland. 2011. “Nkx6 Genes Pattern the Frog Neural Plate and Nkx6.1 Is Necessary for Motoneuron Axon Projection.” Developmental Biology 349 (2): 378–86. 10.1016/j.ydbio.2010.10.030.

Dusabimana, Alfred, Solomon Tsebeni Wafula, Stephen Jada Raimon, et al. 2021. “Effect of Ivermectin Treatment on the Frequency of Seizures in Persons with Epilepsy Infected with Onchocerca Volvulus.” Pathogens 10 (1): 1. 10.3390/pathogens10010021.

Dutta, Suchetana B., Gerit Arne Linneweber, Maheva Andriatsilavo, Peter Robin Hiesinger, and Bassem A. Hassan. 2023. “EGFR-Dependent Suppression of Synaptic Autophagy Is Required for Neuronal Circuit Development.” Current Biology 33 (3): 517–532.e5. 10.1016/j.cub.2022.12.039.

Eberhardt, Jerome, Diogo Santos-Martins, Andreas F. Tillack, and Stefano Forli. 2021. “AutoDock Vina 1.2.0: New Docking Methods, Expanded Force Field, and Python Bindings.” Journal of Chemical Information and Modeling 61 (8): 3891–98. 10.1021/acs.jcim.1c00203.

Estrada-Mondragon, Argel, and Joseph W. Lynch. 2015. “Functional Characterization of Ivermectin Binding Sites in α1β2γ2L GABA(A) Receptors.” Frontiers in Molecular Neuroscience 8 (September). 10.3389/fnmol.2015.00055.

Eswar, Narayanan, Ben Webb, Marc A. Marti Renom, et al. 2006. “Comparative Protein Structure Modeling Using Modeller.” Current Protocols in Bioinformatics 15 (1). 10.1002/0471250953.bi0506s15.

Fu, Wen-Mei. 1995. “Regulatory Role of ATP at Developing Neuromuscular Junctions.” Progress in Neurobiology 47 (1): 31–44. 10.1016/0301-0082(95)00019-R.

GabAllh, Mahmoud, AbdEl-baset El-mashad, Aziza Amin, and Marwa Darweish. 2017. “Pathological Studies on Effects of Ivermectin on Male and Female Rabbits.” Benha Veterinary Medical Journal 32 (1): 104–12. 10.21608/bvmj.2017.31162.

Goyal, Raman, Patricio A. Castro, Jacqueline B. Levin, et al. 2026. “Vesicular Glutamate Release Is Necessary for Neural Tube Formation.” *The Journal of Neuroscience*, February 18, e0681252026. 10.1523/JNEUROSCI.0681-25.2026.

Hattori, M., and E. Gouaux. 2012. “Crystal Structure of the ATP-Gated P2X4 Ion Channel in the ATP-Bound, Open State at 2.8 Angstroms: 4dw1.” April 25. 10.2210/pdb4dw1/pdb.

Hinsby, Anders, M. 2004. “Molecular Mechanisms of NCAM Function.” Frontiers in Bioscience 9 (1–3): 2227. 10.2741/1393.

Huang, Xinying, Junjian Jiang, and Jianguang Xu. 2022. “Denervation-Related Neuromuscular Junction Changes: From Degeneration to Regeneration.” Frontiers in Molecular Neuroscience 14 (February): 810919. 10.3389/fnmol.2021.810919.

Illes, Peter, Christa E. Müller, Kenneth A. Jacobson, et al. 2021. “Update of P2X Receptor Properties and Their Pharmacology: IUPHAR Review 30.” British Journal of Pharmacology 178 (3): 489–514. 10.1111/bph.15299.

Jamieson, D., and A. Roberts. 2000. “Responses of Young Xenopus Laevis Tadpoles to Light Dimming: Possible Roles for the Pineal Eye.” The Journal of Experimental Biology 203 (Pt 12): 1857–67. 10.1242/jeb.203.12.1857.

Johnston, Graham Ar. 2013. “Advantages of an Antagonist: Bicuculline and Other GABA Antagonists.” British Journal of Pharmacology 169 (2): 328–36. 10.1111/bph.12127.

Jones, Emilie, Jay Miguel Fonticella, and Kelly A. McLaughlin. 2025. “Identification and Characterization of Static Craniofacial Defects in Pre-Metamorphic Xenopus Laevis Tadpoles.” Journal of Developmental Biology 13 (3): 26. 10.3390/jdb13030026.

Kaeser, Gwendolyn E., Brian A. Rabe, and Margaret S. Saha. 2011. “Cloning and Characterization of GABA_A_ α Subunits and GABA_B_ Subunits in *Xenopus Laevis* during Development.” Developmental Dynamics 240 (4): 862–73. 10.1002/dvdy.22580.

Khakh, Baljit S., and R. Alan North. 2012. “Neuromodulation by Extracellular ATP and P2X Receptors in the CNS.” Neuron 76 (1): 51–69. 10.1016/j.neuron.2012.09.024.

Khakh, Baljit S., William R. Proctor, Thomas V. Dunwiddie, Cesar Labarca, and Henry A. Lester. 1999. “Allosteric Control of Gating and Kinetics at P2X_4_Receptor Channels.” The Journal of Neuroscience 19 (17): 7289–99. 10.1523/jneurosci.19-17-07289.1999.

Krause, Ryoko M., Bruno Buisson, Sonia Bertrand, et al. 1998. “Ivermectin: A Positive Allosteric Effector of the Α7 Neuronal Nicotinic Acetylcholine Receptor.” Molecular Pharmacology 53 (2): 283–94. 10.1124/mol.53.2.283.

Krůšek, Jan, and Hana Zemková. 1994. “Effect of Ivermectin on γ-Aminobutyric Acid-Induced Chloride Currents in Mouse Hippocampal Embryonic Neurones.” European Journal of Pharmacology 259 (2): 121–28. 10.1016/0014-2999(94)90500-2.

Latapiat, Verónica, Felipe E. Rodríguez, Francisca Godoy, Felipe A. Montenegro, Nelson P. Barrera, and Juan P. Huidobro-Toro. 2017. “P2X4 Receptor in Silico and Electrophysiological Approaches Reveal Insights of Ivermectin and Zinc Allosteric Modulation.” Frontiers in Pharmacology 8 (December): 918. 10.3389/fphar.2017.00918.

Lobikin, Maria, Jean-François Paré, David L. Kaplan, and Michael Levin. 2015. “Selective Depolarization of Transmembrane Potential Alters Muscle Patterning and Muscle Cell Localization in Xenopus Laevis Embryos.” The International Journal of Developmental Biology 59 (7-8–9): 303–11. 10.1387/ijdb.150198ml.

Löscher, Wolfgang. 2023. “Is the Antiparasitic Drug Ivermectin a Suitable Candidate for the Treatment of Epilepsy?” Epilepsia 64 (3): 553–66. 10.1111/epi.17511.

Mackay, Laurent, Hana Zemkova, Stanko S. Stojilkovic, Arthur Sherman, and Anmar Khadra. 2017. “Deciphering the Regulation of P2X4 Receptor Channel Gating by Ivermectin Using Markov Models.” PLOS Computational Biology 13 (7): e1005643. 10.1371/journal.pcbi.1005643.

Mandro, Michel, Joseph Nelson Siewe Fodjo, Alfred Dusabimana, et al. 2020. “Single versus Multiple Dose Ivermectin Regimen in Onchocerciasis-Infected Persons with Epilepsy Treated with Phenobarbital: A Randomized Clinical Trial in the Democratic Republic of Congo.” Pathogens 9 (3): 205. 10.3390/pathogens9030205.

Messenger, N. J., S. J. Rowe, and A. E. Warner. 1999. “The Neurotransmitter Noradrenaline Drives*Noggin*-Expressing Ectoderm Cells to Activate*N-Tubulin*and Become Neurons.” Developmental Biology 205 (2): 224–32. 10.1006/dbio.1998.9125.

Millar-Obreque, Camila, Vicente González-Muñoz, Ana M. Marileo, et al. 2025. “In Silico Characterization of Gelsemium Compounds as Glycine Receptor Ligands.” Compounds 5 (4): 40. 10.3390/compounds5040040.

Moiseiwitsch, Julian R. D. 2000. “The Role of Serotonin and Neurotransmitters During Craniofacial Development.” Critical Reviews in Oral Biology & Medicine 11 (2): 230–39. 10.1177/10454411000110020601.

Moreau, Marc, Isabelle Néant, Sarah E. Webb, Andrew L. Miller, Jean-François Riou, and Catherine Leclerc. 2016. “Ca2+ Coding and Decoding Strategies for the Specification of Neural and Renal Precursor Cells during Development.” Cell Calcium 59 (2–3): 75–83. 10.1016/j.ceca.2015.12.003.

Nguyen, Nguyen Thanh, Trung Hai Nguyen, T. Ngoc Han Pham, et al. 2020. “Autodock Vina Adopts More Accurate Binding Poses but Autodock4 Forms Better Binding Affinity.” Journal of Chemical Information and Modeling 60 (1): 204–11. 10.1021/acs.jcim.9b00778.

Nicolas, Patricia, Marta F. Maia, Quique Bassat, et al. 2020. “Safety of Oral Ivermectin during Pregnancy: A Systematic Review and Meta-Analysis.” The Lancet Global Health 8 (1): e92–100. 10.1016/S2214-109X(19)30453-X.

Nieuwkoop, Pieter Dirk, Jacob Faber, John Gerhart, and Marc Kirschner. 1994. Normal Table of Xenopus Laevis (Daudin): A Systematical and Chronological Survey of the Development from the Fertilized Egg till the End of Metamorphosis. Facsim. ed. With Hubrecht laboratorium. Garland.

O’Rahilly, Ronan, and Fabiola Müller. 2007. “Neurulation in the Normal Human Embryo.” In Novartis Foundation Symposia, 1st ed., edited by Gregory Bock and Joan Marsh. Wiley. 10.1002/9780470514559.ch5.

Parisi, Débora P., Satiro A. R. Santos, Danilo Cabral, et al. 2019. “Therapeutical Doses of Ivermectin and Its Association with Stress Disrupt Motor and Social Behaviors of Juvenile Rats and Serotonergic and Dopaminergic Systems.” Research in Veterinary Science 124 (June): 149–57. 10.1016/j.rvsc.2019.03.009.

Paudel, Sudip, Regan Sindelar, and Margaret Saha. 2018. “Calcium Signaling in Vertebrate Development and Its Role in Disease.” International Journal of Molecular Sciences 19 (11): 3390. 10.3390/ijms19113390.

Phelps, Patricia E., Robert P. Barber, Lynn A. Brennan, Victor M. Maines, Paul M. Salvaterra, and James E. Vaughn. 1990. “Embryonic Development of Four Different Subsets of Cholinergic Neurons in Rat Cervical Spinal Cord.” Journal of Comparative Neurology 291 (1): 9–26. 10.1002/cne.902910103.

Phillis, John W. 2024. Adenosine and Adenine Nucleotides As Regulators of Cellular Function. CRC Press. 10.1201/9781003574996.

Pohanka, Miroslav. 2012. “Alpha7 Nicotinic Acetylcholine Receptor Is a Target in Pharmacology and Toxicology.” International Journal of Molecular Sciences 13 (2): 2219–38. 10.3390/ijms13022219.

Priel, Avi, and Shai D. Silberberg. 2004. “Mechanism of Ivermectin Facilitation of Human P2X4 Receptor Channels.” The Journal of General Physiology 123 (3): 281–93. 10.1085/jgp.200308986.

Roberts. 2010. “How Neurons Generate Behaviour in a Hatchling Amphibian Tadpole: An Outline.” Frontiers in Behavioral Neuroscience, ahead of print. 10.3389/fnbeh.2010.00016.

Roberts, A. 1990. “How Does a Nervous System Produce Behaviour? A Case Study in Neurobiology.” Science Progress 74 (293 Pt 1): 31–51.

Rodrigues, Ricardo J., Joana M. Marques, and Rodrigo A. Cunha. 2019. “Purinergic Signalling and Brain Development.” *Seminars in Cell & Developmental Biology*, Mechanisms of neural differentiation and integration, vol. 95 (November): 34–41. 10.1016/j.semcdb.2018.12.001.

Saito, Kenzi, Toshikazu Kakizaki, Ryotaro Hayashi, et al. 2010. “The Physiological Roles of Vesicular GABA Transporter during Embryonic Development: A Study Using Knockout Mice.” Molecular Brain 3 (1): 40. 10.1186/1756-6606-3-40.

Saul, Anika, Ralf Hausmann, Achim Kless, and Annette Nicke. 2013. “Heteromeric Assembly of P2X Subunits.” Frontiers in Cellular Neuroscience 7. 10.3389/fncel.2013.00250.

Schindelin, Johannes, Ignacio Arganda-Carreras, Erwin Frise, et al. 2012. “Fiji: An Open-Source Platform for Biological-Image Analysis.” Nature Methods 9 (7): 676–82. 10.1038/nmeth.2019.

Sequerra, Eduardo B., Raman Goyal, Patricio A. Castro, Jacqueline B. Levin, and Laura N. Borodinsky. 2018. “NMDA Receptor Signaling Is Important for Neural Tube Formation and for Preventing Antiepileptic Drug-Induced Neural Tube Defects.” The Journal of Neuroscience 38 (20): 4762–73. 10.1523/jneurosci.2634-17.2018.

Shaker, Mohammed R., Ju-Hyun Lee, and Woong Sun. 2021. “Embryonal Neuromesodermal Progenitors for Caudal Central Nervous System and Tissue Development.” Journal of Korean Neurosurgical Society 64 (3): 359–66. 10.3340/jkns.2020.0359.

Shan, Qiang, Justine L. Haddrill, and Joseph W. Lynch. 2001. “Ivermectin, an Unconventional Agonist of the Glycine Receptor Chloride Channel.” Journal of Biological Chemistry 276 (16): 12556–64. 10.1074/jbc.M011264200.

Shen, Cheng, Yuqing Zhang, Wenwen Cui, et al. 2023. “Structural Insights into the Allosteric Inhibition of P2X4 Receptors.” Nature Communications 14 (1): 6437. 10.1038/s41467-023-42164-y.

Shen, Min□yi, and Andrej Sali. 2006. “Statistical Potential for Assessment and Prediction of Protein Structures.” Protein Science 15 (11): 2507–24. 10.1110/ps.062416606.

Siewe Fodjo, Joseph Nelson, Michel Mandro, Deby Mukendi, et al. 2019. “Onchocerciasis-Associated Epilepsy in the Democratic Republic of Congo: Clinical Description and Relationship with Microfilarial Density.” PLOS Neglected Tropical Diseases 13 (7): e0007300. 10.1371/journal.pntd.0007300.

Sim, Joan A., Séverine Chaumont, Jihoon Jo, et al. 2006. “Altered Hippocampal Synaptic Potentiation in P2X_4_Knock-Out Mice.” The Journal of Neuroscience 26 (35): 9006–9. 10.1523/jneurosci.2370-06.2006.

Slusarski, Diane C., and Francisco Pelegri. 2007. “Calcium Signaling in Vertebrate Embryonic Patterning and Morphogenesis.” Developmental Biology 307 (1): 1–13. 10.1016/j.ydbio.2007.04.043.

Spencer, Kira A., Yesser Hadj Belgacem, Olesya Visina, Sangwoo Shim, Henry Genus, and Laura N. Borodinsky. 2019. “Growth at Cold Temperature Increases the Number of Motor Neurons to Optimize Locomotor Function.” Current Biology 29 (11): 1787–1799.e5. 10.1016/j.cub.2019.04.072.

Spuy, Marise van der, Jian Xiong Wang, Dagmara Kociszewska, and Melanie D. White. 2023. “The Cellular Dynamics of Neural Tube Formation.” Biochemical Society Transactions 51 (1): 343–52. 10.1042/BST20220871.

Steventon, Ben, Claudio Araya, Claudia Linker, Sei Kuriyama, and Roberto Mayor. 2009. “Differential Requirements of BMP and Wnt Signalling during Gastrulation and Neurulation Define Two Steps in Neural Crest Induction.” Development 136 (5): 771–79. 10.1242/dev.029017.

Stokes, Leanne, Janice A. Layhadi, Lucka Bibic, Kshitija Dhuna, and Samuel J. Fountain. 2017. “P2X4 Receptor Function in the Nervous System and Current Breakthroughs in Pharmacology.” Frontiers in Pharmacology 8 (May): 291. 10.3389/fphar.2017.00291.

Sullivan, Kelly G., and Michael Levin. 2016. “Neurotransmitter Signaling Pathways Required for Normal Development in Xenopus Laevis Embryos: A Pharmacological Survey Screen.” Journal of Anatomy 229 (3): 483–502. 10.1111/joa.12467.

Suyama, S., T. Sunabori, H. Kanki, et al. 2012. “Purinergic Signaling Promotes Proliferation of Adult Mouse Subventricular Zone Cells.” Journal of Neuroscience 32 (27): 9238–47. 10.1523/JNEUROSCI.4001-11.2012.

Suzuki, Makoto, Masanao Sato, Hiroshi Koyama, et al. 2017. “Distinct Intracellular Ca2+ Dynamics Regulate Apical Constriction and Differentially Contribute to Neural Tube Closure.” Development 144 (7): 1307–16. 10.1242/dev.141952.

Ten Donkelaar, Hans J., Martin Lammens, and Akira Hori. 2006. Clinical Neuroembryology. Springer Berlin Heidelberg. 10.1007/3-540-34659-7.

Tiboni, Gian Mario, Adalisa Ponzano, Alessio Ferrone, Sara Franceschelli, Lorenza Speranza, and Antonia Patruno. 2021. “Valproic Acid Alters Nitric Oxide Status in Neurulating Mouse Embryos.” Reproductive Toxicology 99 (January): 152–59. 10.1016/j.reprotox.2020.08.012.

Tovar, Lina Mariana, Carlos Felipe Burgos, Gonzalo E. Yévenes, et al. 2023. “Understanding the Role of ATP Release through Connexins Hemichannels during Neurulation.” International Journal of Molecular Sciences 24 (3): 2159. 10.3390/ijms24032159.

Trott, Oleg, and Arthur J. Olson. 2010. “AutoDock Vina: Improving the Speed and Accuracy of Docking with a New Scoring Function, Efficient Optimization, and Multithreading.” Journal of Computational Chemistry 31 (2): 455–61. 10.1002/jcc.21334.

Ulrich, Henning, and Peter Illes. 2014. “P2X Receptors in Maintenance and Differentiation of Neural Progenitor Cells.” Neural Regeneration Research 9 (23): 2040. 10.4103/1673-5374.147925.

Vijayraghavan, Deepthi S., and Lance A. Davidson. 2017. “Mechanics of Neurulation: From Classical to Current Perspectives on the Physical Mechanics That Shape, Fold, and Form the Neural Tube.” Birth Defects Research 109 (2): 153–68. 10.1002/bdra.23557.

Von Kügelgen, Ivar. 2011. “Molecular Pharmacology, Physiology, and Structure of the P2Y Receptors.” In Advances in Pharmacology, by T. Kendall Harden. Elsevier. 10.1016/b978-0-12-385526-8.00012-6.

Wallace, Andrew C., Roman A. Laskowski, and Janet M. Thornton. 1995. “LIGPLOT: A Program to Generate Schematic Diagrams of Protein-Ligand Interactions.” *Protein Engineering*, Design and Selection 8 (2): 127–34. 10.1093/protein/8.2.127.

Webb, Sarah E., and Andrew L. Miller. 2003. “Calcium Signalling during Embryonic Development.” Nature Reviews Molecular Cell Biology 4 (7): 539–51. 10.1038/nrm1149.

Werner, Stefan, Stefanie Mesch, Roman C. Hillig, et al. 2019. “Discovery and Characterization of the Potent and Selective P2X4 Inhibitor *N*-[4-(3-Chlorophenoxy)-3-Sulfamoylphenyl]-2-Phenylacetamide (BAY-1797) and Structure-Guided Amelioration of Its CYP3A4 Induction Profile.” Journal of Medicinal Chemistry 62 (24): 11194–217. 10.1021/acs.jmedchem.9b01304.

Whitaker, Michael. 2006. “Calcium at Fertilization and in Early Development.” Physiological Reviews 86 (1): 25–88. 10.1152/physrev.00023.2005.

Zaganjor, Ibrahim, Ahlia Sekkarie, Becky L. Tsang, et al. 2016. “Describing the Prevalence of Neural Tube Defects Worldwide: A Systematic Literature Review.” PLOS ONE 11 (4): e0151586. 10.1371/journal.pone.0151586.

Zimmermann, Herbert. 2011. “Purinergic Signaling in Neural Development.” *Seminars in Cell & Developmental Biology*, Recent Progress in Ion Channels, vol. 22 (2): 194–204. 10.1016/j.semcdb.2011.02.007.

Zlotos, Darius P., Yasmine M. Mandour, and Anders A. Jensen. 2022. “Strychnine and Its Mono-and Dimeric Analogues: A Pharmaco-Chemical Perspective.” Natural Product Reports 39 (10): 1910–37. 10.1039/D1NP00079A.

Zschätzsch, Marlen, Carlos Oliva, Marion Langen, et al. 2014. “Regulation of Branching Dynamics by Axon-Intrinsic Asymmetries in Tyrosine Kinase Receptor Signaling.” eLife 3 (April): e01699. 10.7554/eLife.01699.

