## Supplemental Material for "Ivermectin exposition during neurulation induces Neural tube defects and neuromuscular alterations in *Xenopus laevis* through purinergic P2X4-signaling"

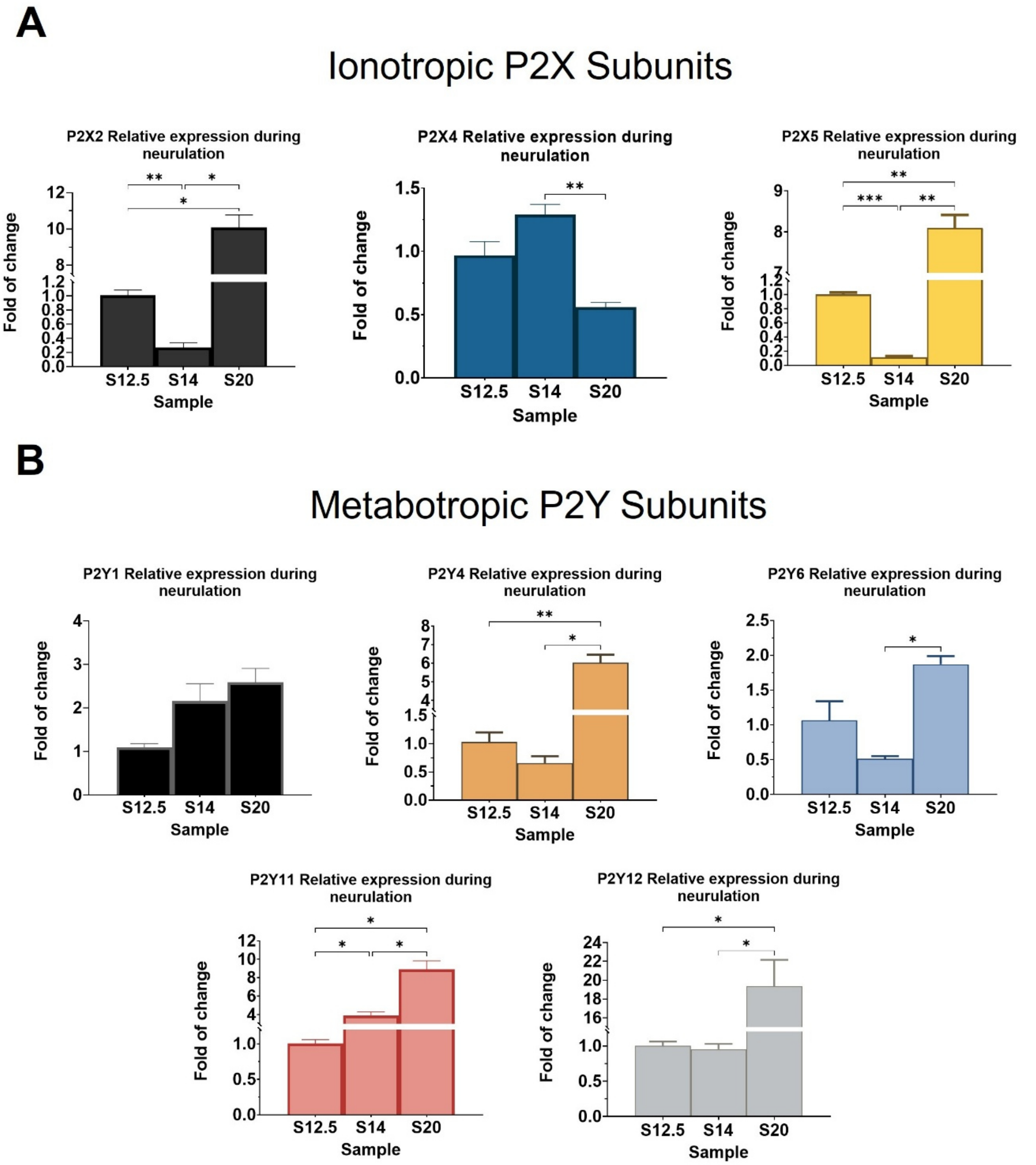


Supplementary figure 1. **RT-qPCR screening analysis of purinergic subunits during neurulation.** (A) Fold of change of ionotropic P2X2,4,5 mRNA subunits normalized to stage 12.5. (B) Fold of change of metabotropic P2Y1,4,6,11,12 mRNA subunits normalized to stage S12.5. N=3. Brown-Forsythe and Welch ANOVA, Dunnet's correction, (* P<0.05, ** P<0.01, *** P<0.001). The error bars of the data correspond to the standard error of the mean (SEM)

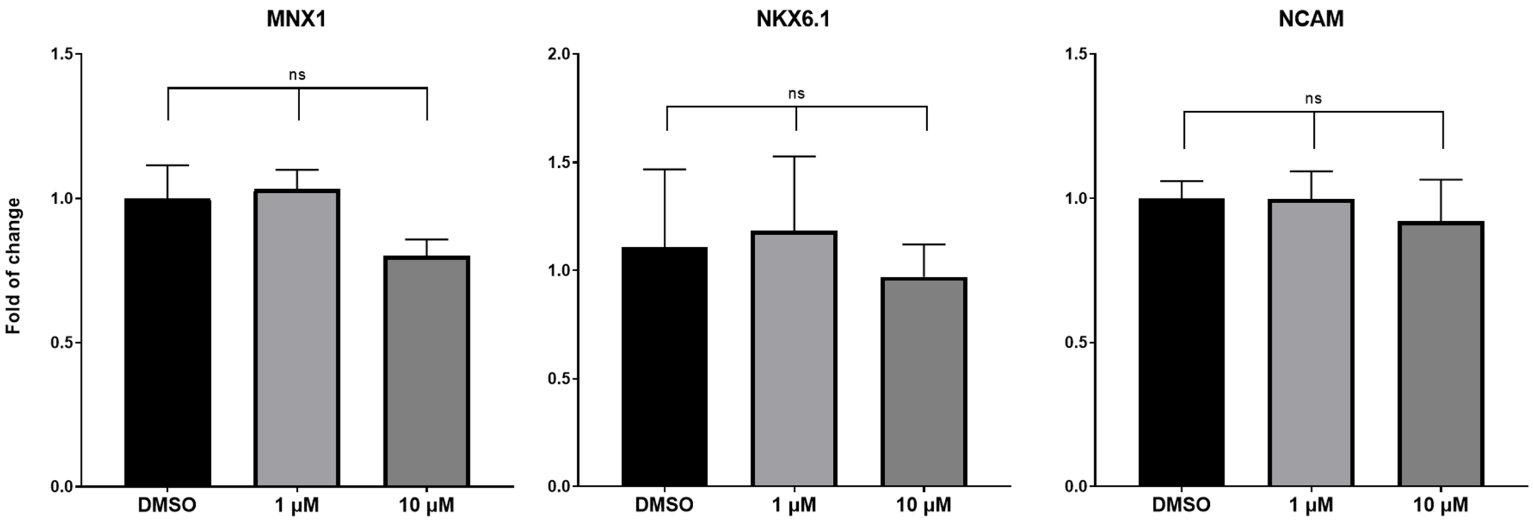


Supplementary figure 2. **Relative expression of motoneurons related transcription factors at tadpole stage 45.** Fold of change of NKX 6.1, NCAM and MNX1 transcripts of IVM treated conditions (1 and 10 µM) compared with control (DMSO) through RT-qPCR assays. N=3. One-way ANOVA, Dunnet's correction, (ns=not significative). The error bars of the data correspond to the standard error of the mean (SEM).

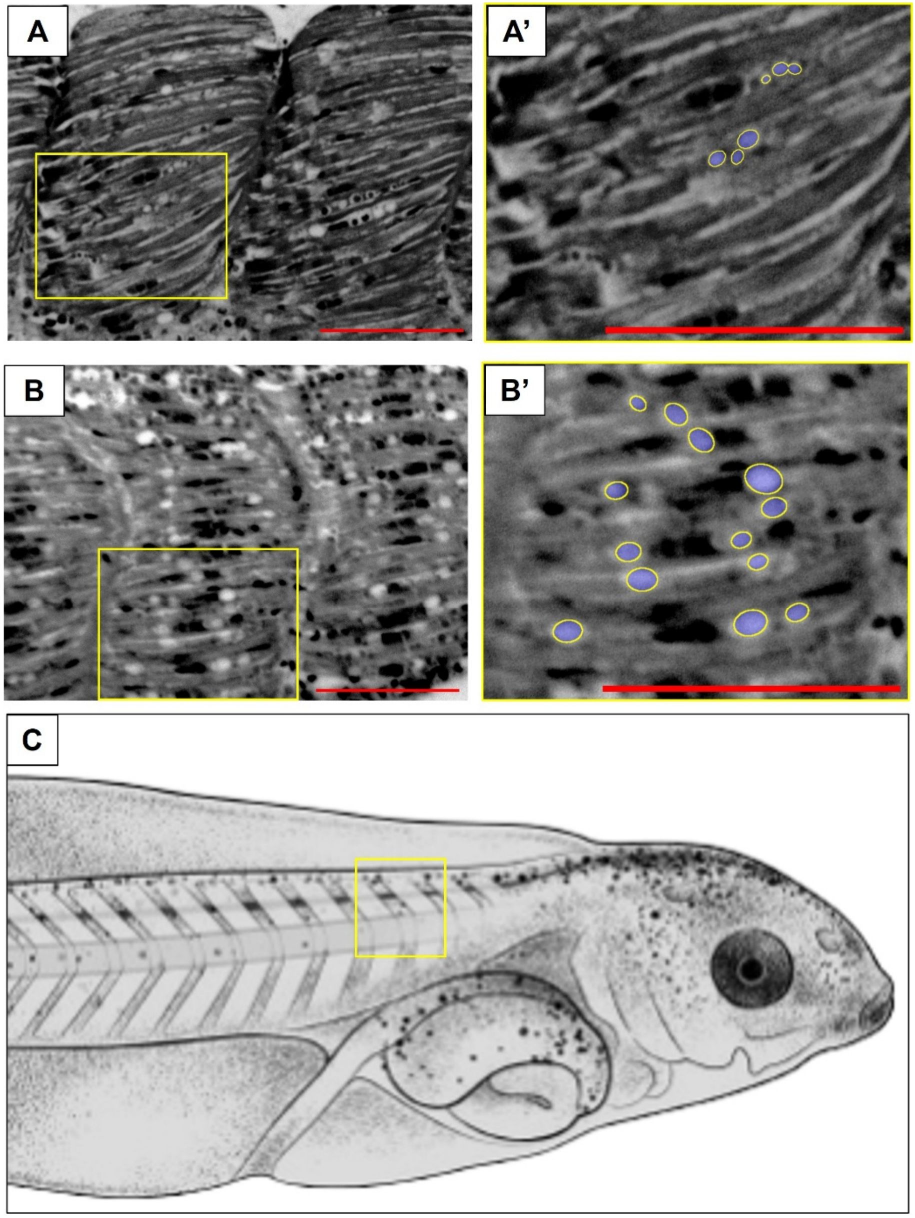


Supplementary figure 3. **Transversal hematoxylin & eosin muscle sections of chevron 4-5 of controls and neurula-IVM treatment of stage 45 tadpoles. (A)** Typical pattern of horizontal muscle fibers distribution on control tadpoles. **(A')** Zoom on vacuolized presented on control section. **(B).** Representative Muscle fibers organization on neurula-IVM treated (10 µM) tadpoles. **(B')** Zoom of area showing an altered number of muscle fibers vacuolized presented on 10 µM IVM treatment tadpoles. **(C)** Ilustrated tadpole with yellow box indicating the region shown on A, A', B, B' scale bar:100 µm.

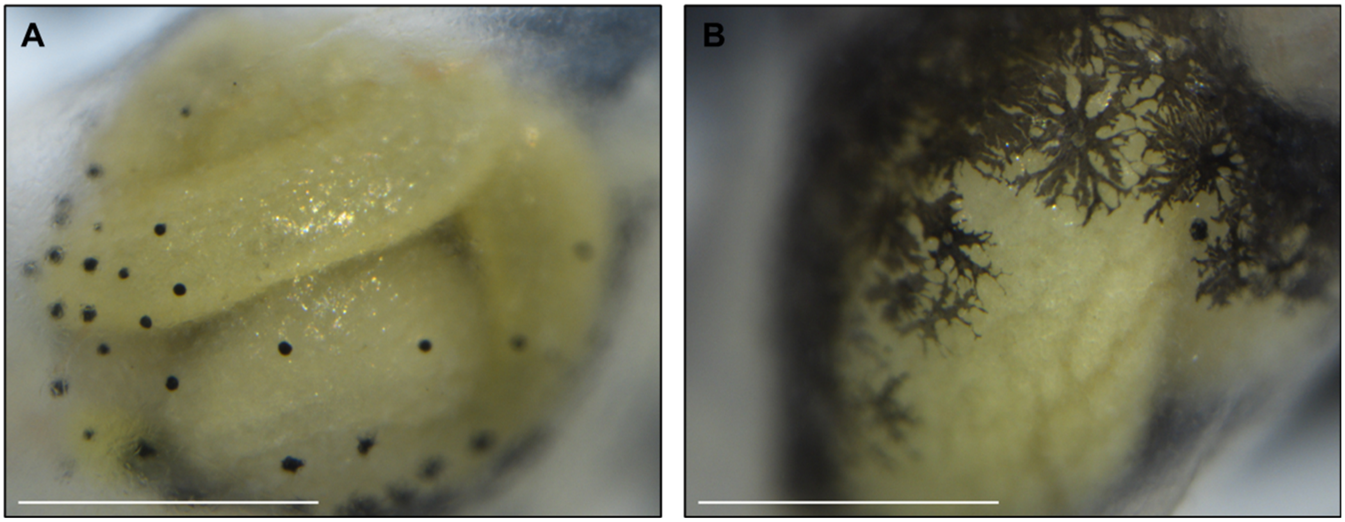


Supplementary figure 4. **Effect of 100 µM ivermectin on digestive system of stage 45 Xenopus tadpoles. (A)** Ventral view of consolidated digestive system and digestive tube in control tadpoles. Typical normal melanocyte phenotype (dark cells) is presented **(B)** Ventral view of alter digestive system on 100 µM IVM neurula-treated tadpoles, in which the digestive tube cannot be identified. Hyper arborized melanocyte phenotype (dark star shape cells) is presented. A, B scale bar: 100 µm.

| Transcription factors | |
|---|---|
| Subunit | Sequence (5'-3') |
| mnx1.L F | TTCTGACTGCACATCCGAGG |
| mnx1.L R | GGGTGATACTGGCTCAGTGG |
| ncam1.L F | ACAGATGTCAAGACCCC |
| ncam1.L R | ATTCCCAATAACACCACCCCT |
| nkx6-1.L F | CCGGAAGGACTCCCATCTTT |
| nkx6-1.L R | AACACAGAGCCCTGATTGGG |

**Supplementary table 1. Primer sequences used in stage 45 tadpole.** R=Reverse primer, F=Forward primer, S=Short, L=Long.
